# SWI/SNF-Dependent Chromatin Remodelling Establishes Gene Regulatory Programs in Early Human Embryos and Blastoids

**DOI:** 10.64898/2025.12.01.691499

**Authors:** Sam S.F.A. van Knippenberg, Maria Tryfonos, Joke De Busscher, Elena Biondi, Mairim Alexandra Solis, Chloé Lorent, Océane Girard, Suresh Poovathingal, Marta Wojno, Sherif Khodeer, Jade De Clercq, Antonina Mikorska, Inge Smeers, Eva Wigerinck, Yara Meynen, Thierry Voet, Laurent David, Hilde Van de Velde, Vincent Pasque

## Abstract

Establishment of cell lineages during development is regulated by transcription factors binding at cis-regulatory elements to activate gene regulatory programs. Many transcription factors require chromatin remodellers for accessibility at their target sites. Although these key principles of gene regulation in mammalian development have emerged, the contribution of chromatin remodellers to early human embryogenesis remains unknown. Here, we show that the SWI/SNF ATPases SMARCA2 and SMARCA4 are required for establishing the epiblast and trophectoderm fates during human pre-implantation development and facilitate accessibility at regulatory elements. We find that degradation of SWI/SNF ATPases disrupts epiblast formation in blastoids and enhances trophectoderm specification, while also showing transcriptional and chromatin misregulation in TE-like cells. In human embryos, SWI/SNF perturbation impaired blastocyst formation and the establishment of the inner cell mass. Single-nucleus chromatin accessibility and transcriptome profiling in blastoids reveals that the SWI/SNF complex safeguards the naïve epiblast and trophectoderm programs and facilitates enhancer and transcription factor motif accessibility. These findings identify SWI/SNF chromatin remodellers as critical regulators of embryonic lineage specification during human pre-implantation development.

## Introduction

Human embryogenesis begins with the fertilisation of the oocyte by a spermatozoon, resulting in the formation of the totipotent zygote. The zygote subsequently undergoes cleavage divisions, forming blastomeres. The blastomeres compact, forming the morula, which cavitates to form a blastocyst around 5 days post-fertilisation (dpf)^1^. Around this time, the first lineage segregation occurs, where the embryo gradually diverges into the extraembryonic trophectoderm (TE) and the inner cell mass (ICM)^1^. Subsequently, the second lineage segregation involves the ICM diverging into the embryonic epiblast (EPI) and the extraembryonic hypoblast (HYPO)^2,3^.

Cells of the early human blastocyst possess a broad lineage potential^4^. This broad lineage potential is maintained in naïve human pluripotent stem cells (hPSCs), which resemble the cells of the pre-implantation EPI. Naïve hPSCs can differentiate into the three founding lineages of the human embryo, including embryonic EPI, extraembryonic TE and HYPO lineages in adherent cultures^5–10^ and in 3D differentiation systems^11–16^. Notably, naïve hPSCs can self-assemble into blastocyst-like structures termed blastoids^11–15^, which resemble the human blastocyst at a transcriptional and morphological level^11–14,17^.

Blastoids, and other stem cell-based embryo models, provide a promising platform to study the gene regulatory mechanisms underlying early human embryogenesis *in vitro*^18^. However, although human blastoids have been shown to model HYPO specification^19^, the extent to which they can recapitulate human embryonic TE specification remains unclear.

The establishment of cell fates is regulated by transcription factor (TF)-driven gene regulatory programs^20–24^. Gene regulatory programs are specific per cell type and facilitated by chromatin remodelling complexes, which enable chromatin accessibility at the correct cis-regulatory elements^25–28^. The establishment of cell fates during early development is accompanied by drastic changes in the chromatin landscape^27^. These changes in the chromatin landscape are regulated by chromatin remodelling complexes, among which is the SWItch/Sucrose Non-Fermentable (SWI/SNF) complex, also known as the Brahma Associated Factor (BAF) complex^29^. How chromatin remodelling by the SWI/SNF complex impacts gene regulatory programs in early human embryonic development remains largely unknown.

The SWI/SNF complex is an ATP-dependent chromatin remodelling complex that promotes chromatin accessibility through nucleosome ejection and sliding^30^, which impacts cell identity. The SWI/SNF complex is thought to interact with TFs and RNA polymerase II to regulate enhancer and promoter accessibility at genes related to cell identity, development, and cell signalling, but not necessarily for housekeeping and viability-related genes^31–34^. The ATPases of the SWI/SNF complex are either SMARCA2, also termed Brahma (BRM), or SMARCA4, also known as Brahma-Related Gene 1 (BRG1).

In mice, the SWI/SNF complex has been shown to regulate early lineage specification^35^ and *Smarca4*, but not *Smarca2*, was reported to regulate naïve and primed pluripotency^36,37^ and pre-implantation development^38^. Perturbation of the SWI/SNF complex in murine pre-implantation embryos leads to ectopic expression of pluripotency factors in TE cells, and a failure to form outgrowths from both TE and ICM cells^35,36,39,40^. Interestingly, although *Smarca2* is dispensable for murine development^41^, *Smarca2* can compensate for *Smarca4* haploinsufficiency^38^. These data suggest that the correct dosage of SWI/SNF ATPases may be important during mouse development.

Species-specific differences exist between the function of murine and human SWI/SNF complex ATPases in pluripotency^42^. In contrast to mice, both SWI/SNF ATPases, *SMARCA2* and *SMARCA4*, are expressed in naïve hPSCs^42^. Furthermore, human naïve pluripotency is only lost upon the depletion of both ATPases, indicating at least some level of redundancy^37,42^. In naïve hPSCs, it has been shown that both SWI/SNF ATPases interact with OCT4 at enhancers associated with TE genes, stem cell maintenance, blastocyst-, and placental development^42^. Furthermore, SMARCA2 is highly enriched in naïve hPSCs compared to primed hPSCs, the latter resembling the post-implantation EPI^43^. SMARCA4 has been shown to facilitate the exit of naïve pluripotency towards the primed pluripotent state^42,44^. These studies have shown the important role of the SWI/SNF complex in regulating human naïve pluripotency. However, the role of the SWI/SNF complex in the establishment of cell fates during early human pre-implantation embryo development and in blastoids has not yet been characterised.

Here, we aimed to investigate the role of SWI/SNF ATPases in human pre-implantation development and early cell fate establishment in embryos and in stem cell-based embryo models. We characterised and perturbed SWI/SNF ATPases during TE induction from naïve hPSCs, blastoid formation, and in human blastocysts. Finally, we performed single-nucleus multiomic and transcriptomic profiling of SWI/SNF-perturbed blastoids. Our results show that the SWI/SNF complex regulates TE specification and safeguards EPI identity. Altogether, our study provides insight into the role of chromatin remodelling in early human embryonic development.

## Results

### SWI/SNF ATPases show distinct developmental expression patterns

To investigate SWI/SNF ATPases in early human development, we characterised their expression patterns in multiple published transcriptome datasets from naïve hPSC-derived trophectoderm (nTE)^6^, human blastoids^13^, and human embryos^12,17,45–49^ **(Fig. 1a)**. Both SWI/SNF ATPases showed a constant expression pattern during naïve-to-TE transition in both nTE and blastoid differentiation, and *SMARCA2* only decreased in HYPO cells during blastoid formation **(Fig. 1a top, middle)**. In the pre-implantation embryo, *SMARCA4* showed a constant RNA expression level throughout developmental time and cell lineages **(Fig. 1a, bottom)**. In contrast, *SMARCA2* expression was more dynamic, gradually decreasing during morula and early blastocyst formation, and subsequently increasing in both EPI and TE lineages, but not in HYPO cells **(Fig. 1a, bottom)**.

**Fig. 1:**
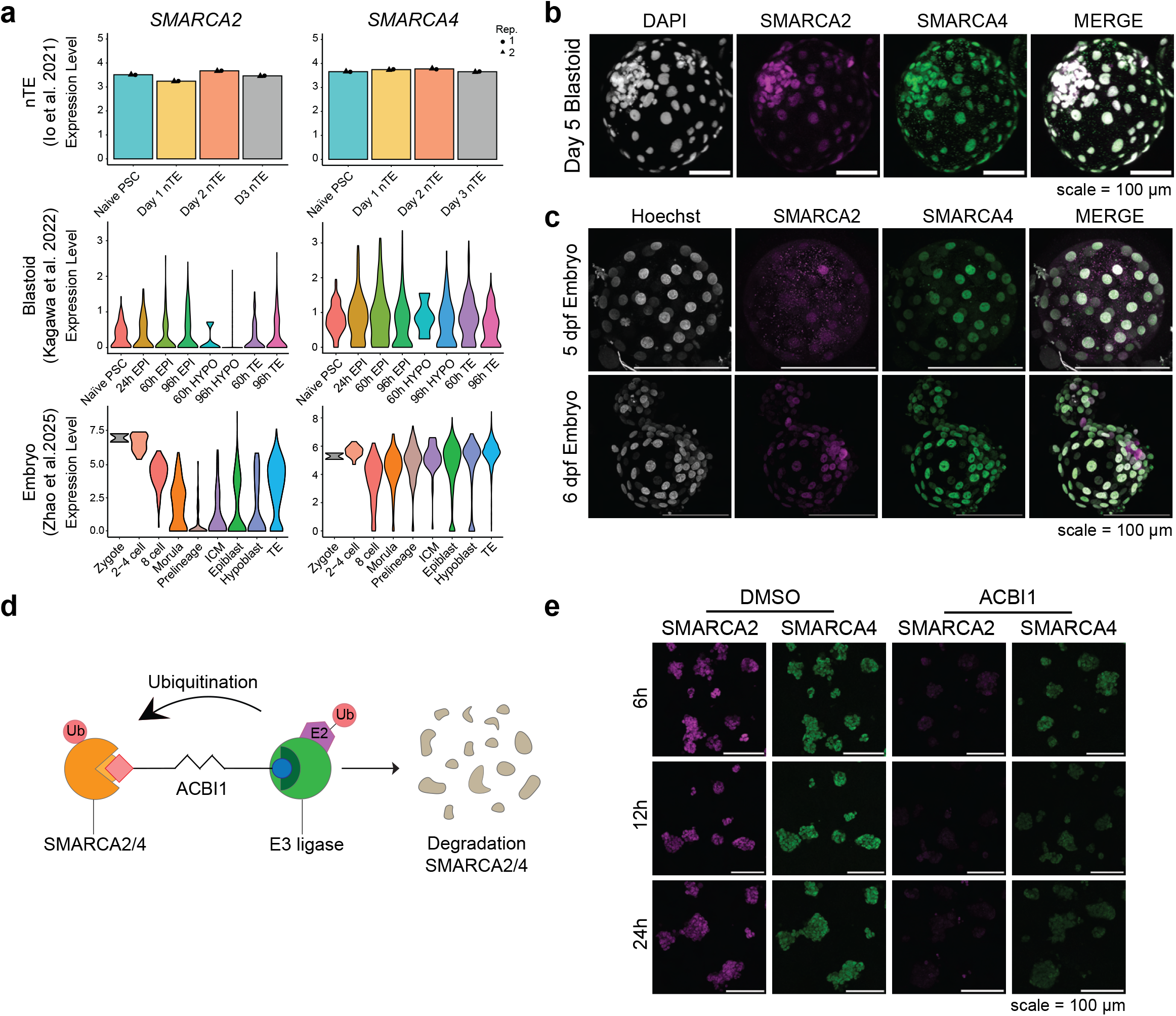
SWI/SNF ATPases show distinct developmental expression patterns. **a** Plots showing gene expression of *SMARCA2* and *SMARCA4* in human naïve hPSC-derived trophectoderm (nTE)^6^ (top), human blastoids^13^ (middle), and human embryos^17^ (bottom). **b, c** Immunofluorescence images of a day 5 human blastoid **(b)** and 5 days post-fertilisation (dpf) and 6 dpf human blastocysts. **c** showing SMARCA2 and SMARCA4 protein expression. Scale bar = 100 µm. N=3 embryos per time point. **d** Schematic representation of the mechanism of action of ACBI1. **e** Immunofluorescence images showing SMARCA2 and SMARCA4 degradation in naïve hPSCs upon 6, 12, and 24 hours of ACBI1 treatment. Scale bar = 100 µm.

Next, we assessed protein expression of SWI/SNF ATPases. We used naïve hPSCs, cultured feeder-free in PXGL^50,51^, nTE cells, differentiated using a protocol from Io et al.^6^, human blastoids, generated using an adapted protocol from Kagawa et al.^13^, and 5 dpf and 6 dpf human supernumerary IVF embryos cultured *ex vivo*. Additionally, we analysed existing proteomic datasets^52,53^. Immunofluorescence analysis showed that, in line with RNA expression, SMARCA4 was present at the protein level in naïve hPSCs, nTE cells and in all cells of human blastoids and blastocysts **(Fig. 1b, c and Supplementary Fig. 1a-d)**. Proteomic analysis revealed that in stem cells, SMARCA4 remains highly expressed in both primed hPSCs and trophoblast stem cells (TSCs), the latter resembling the post-implantation stage of the trophectoderm lineage **(Supplementary Fig. 1e)**. SMARCA4 showed low protein abundance in human pre-implantation embryos^53^ **(Supplementary Fig. 1e)**. Altogether, these data suggest that SMARCA4 protein levels are constant during blastocyst formation and in early human cell lineages.

SMARCA2 exhibited relatively low protein levels in 5 dpf human blastocysts, in line with the gene expression data **(Fig. 1a, bottom, and Fig. 1c)**. However, while gene expression analysis suggested similar expression of *SMARCA2* in TE and EPI cells **(Fig. 1a)**, in adherent cultures, blastoids, and 6 dpf human blastocysts, SMARCA2 was enriched at the protein level in EPI cells compared to TE cells **(Fig. 1b, c and Supplementary Fig. 1a-d)**. Proteomic analysis suggests that SMARCA2 is indeed decreased in primed hPSCs, but that TSCs show a SMARCA2 protein abundance similar to naïve hPSCs **(Supplementary Fig. 1e)**, suggesting that SMARCA2 might be upregulated in later TE lineages. Interestingly, human embryo proteomic analysis suggests that SMARCA2 is highly abundant in the blastocyst stage **(Supplementary Fig. 1e)**. Overall, these data suggest that SWI/SNF ATPases have distinct expression patterns during blastocyst development, where SMARCA4 is consistently present throughout developmental time and lineages, whereas SMARCA2 expression is more dynamic and enriched in EPI cells.

### SWI/SNF ATPase degradation leads to loss of naïve pluripotency

To study the impact of the SWI/SNF complex perturbation on human embryonic development, we aimed to use an inducible perturbation system that can be applied to stem cell lines, blastoids, and *ex vivo* cultured human embryos. As SMARCA2 and SMARCA4 have been shown to have at least some level of redundancy in naïve hPSCs^42^, we opted for a system enabling perturbation of both SWI/SNF ATPases. We used the PROteolysis-TArgeting-Chimaera (PROTAC) ACBI1^54^, enabling fast and specific targeted degradation of both SWI/SNF ATPases through the ubiquitination-proteasome pathway^55^ **(Fig. 1d)**. ACBI1 treatment at 50nM was sufficient for the degradation of both SWI/SNF ATPases within a few hours, with slightly more efficient degradation of SMARCA2 than for SMARCA4 **(Fig. 1e)**. To investigate the effect of SWI/SNF perturbation using the PROTAC on naïve pluripotency, we cultured naïve hPSCs with ACBI1 for 12 days **(Supplementary Fig. 2a)**. We found that naïve hPSCs appeared relatively normal after one passage. However, prolonged ACBI1 exposure induced a gradual decline in cell viability, which prevented the maintenance of naïve hPSCs beyond four passages **(Supplementary Fig. 2b, c)**. These results are consistent with previous findings showing a loss of naïve pluripotency upon combined genetic ablation and knockdown of both SWI/SNF ATPases^42^. However, in contrast to previous findings, we did not observe a collapse of colony morphology. Altogether, these results confirm that SWI/SNF ATPases are essential for the maintenance of naïve human pluripotency.

As SWI/SNF ATPases are required for long-term maintenance of naïve human pluripotency, we next investigated the short-term consequences of SWI/SNF ATPase degradation under naïve conditions. Naïve hPSCs were cultured for three days in either PXGL, supporting naïve pluripotency, or N2B27 basal medium, without additional signalling modulators, and were treated with ACBI1 or DMSO **(Supplementary Fig. 2d)**. In PXGL, cells maintained a dome-shaped colony morphology following three days of ACBI1 treatment, similar to the untreated (WT) and DMSO control conditions **(Supplementary Fig. 2e)**. After three days in N2B27, control conditions contained both dome-shaped and flattened colonies. In contrast, the ACBI1-treated condition displayed a predominantly flattened colony morphology **(Supplementary Fig. 2e)**. Thus, acute SWI/SNF ATPase degradation induces a morphological response upon the removal of naïve pluripotency supporting cues.

Next, we assessed whether this morphological response was accompanied by induction of TE-associated markers. We performed flow cytometry for TE markers TROP2 and ENPEP. ACBI1 treatment modestly increased the levels of both markers in PXGL and N2B27 conditions **(Supplementary Fig. 2f)**. However, immunofluorescence did not show the induction of TE transcription factors GATA3 or TEAD1 following ACBI1 treatment **(Supplementary Fig. 2g)**. Therefore, although acute SWI/SNF ATPase degradation is associated with altered colony morphology and modest upregulation of TE markers, it is insufficient to induce the TE regulatory program under either culture condition.

### SWI/SNF ATPases facilitate naïve hPSC-to-TE conversion

To further investigate the role of the SWI/SNF complex in TE induction, we treated cells with ACBI1 throughout nTE differentiation^6^ **(Supplementary Fig. 3a)**. Under control conditions, cells progressively acquired a TE-like morphology with prominent cell boundaries. This morphological transition was altered in ACBI1-treated cultures **(Supplementary Fig. 3b)**. Quantification of TE-associated surface markers TROP2 and ENPEP by flow cytometry showed nTE conversion efficiencies in WT and DMSO-control conditions that were consistent with previous reports^6^. In contrast, the proportion of TROP2- and ENPEP-positive cells was reduced following ACBI1 treatment, indicating that degradation of SWI/SNF ATPases impairs naïve hPSC-to-TE conversion **(Supplementary Fig. 3c)**.

We next asked whether the reduced frequency of cells expressing TE-associated surface markers was accompanied by altered downregulation of the pluripotency programme and/or impaired acquisition of TE-associated genes. Time-course RT-qPCR showed progressive reduction in expression of the pluripotency-associated genes *POU5F1, NANOG, KLF17,* and *PRDM14* during nTE differentiation **(Supplementary Fig. 3d)**. These transcriptional changes were not detectably changed by ACBI1 treatment **(Supplementary Fig. 3d)**. Consistent with these findings, immunofluorescence analysis showed a similar temporal reduction in NANOG expression in control and ACBI1-treated conditions **(Supplementary Fig. 3e-g).** These data suggest that silencing of the pluripotency program during nTE differentiation is not affected by SWI/SNF perturbation.

Time-course RT-qPCR showed that TE-associated genes *GATA2* and *GATA3* were gradually increased throughout nTE differentiation, as expected **(Supplementary Fig. 3d)**. The induction of TE gene expression was unaffected by ACBI1 treatment **(Supplementary Fig. 3d)**. However, even though *GATA2* and *GATA3* RNA levels were unaffected by ACBI1 treatment, immunofluorescence against GATA2 showed that degradation of SWI/SNF ATPases led to impaired TE induction **(Supplementary Fig. 3e-g)**. These data suggest that the SWI/SNF complex facilitates the acquisition of the TE fate during nTE differentiation.

### The SWI/SNF complex regulates TE induction and EPI formation in human blastoids

Next, we investigated how SWI/SNF ATPase degradation impacts blastoid formation and lineage composition. For this, we generated human blastoids and treated them with ACBI1 to induce degradation of the SWI/SNF ATPases throughout blastoid formation **(Fig. 2a-c)**. We observed that ACBI1-treated blastoids cavitated earlier than the DMSO-treated controls **(Fig. 2d and Supplementary Fig. 4a)**. In line with enhanced cavitation, blastoid growth was accelerated upon SWI/SNF perturbation **(Fig. 2e)**. These observations indicate that the SWI/SNF complex regulates human blastoid formation.

**Fig. 2:**
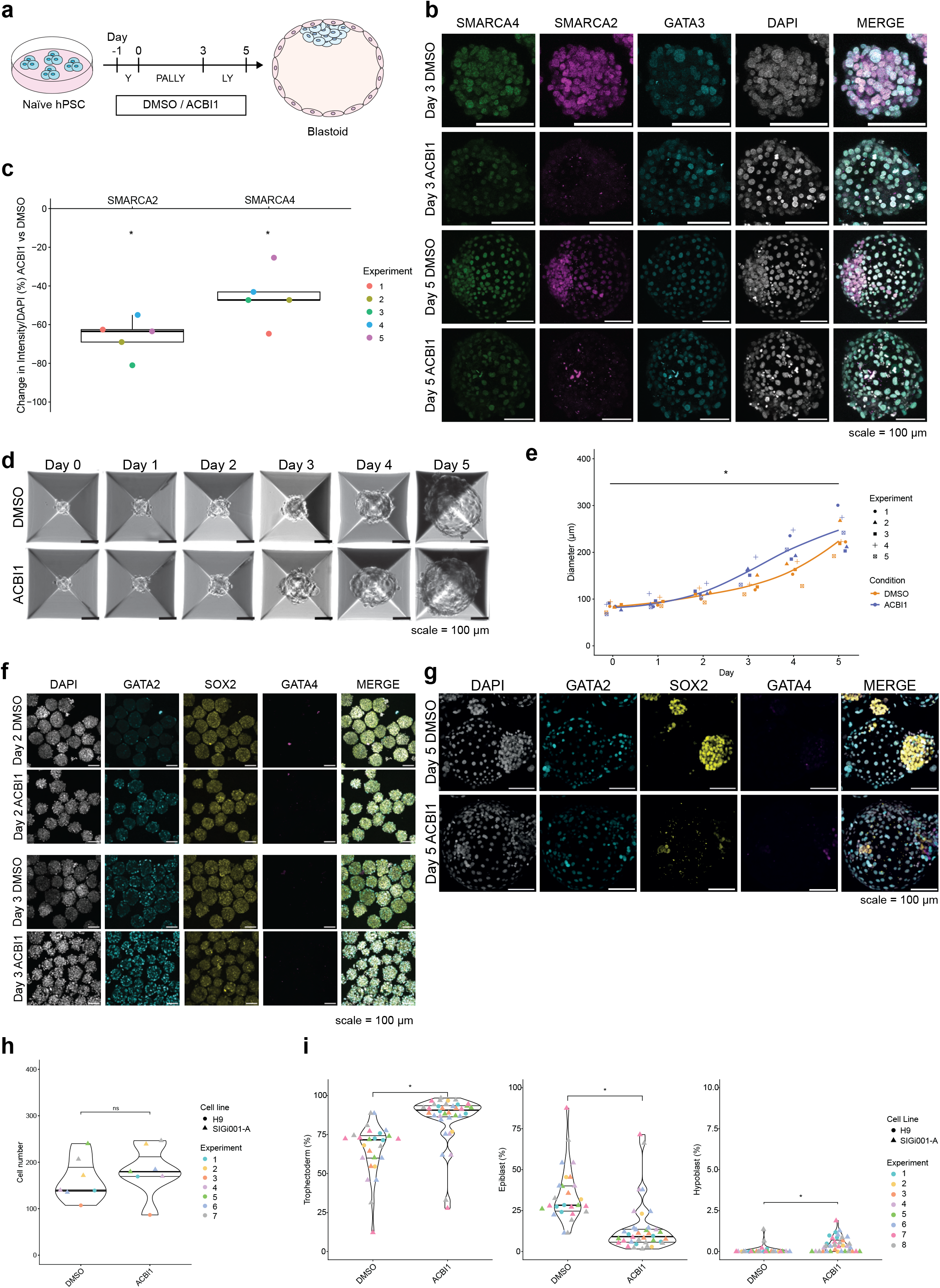
SWI/SNF ATPases regulate trophectoderm specification and safeguard epiblast formation in blastoids. **a** Schematic overview of the experimental set-up of 50 nM ACBI1 treatment during blastoid formation. **b** Immunofluorescence images showing GATA3 protein expression and SMARCA2 and SMARCA4 degradation in day 3 and day 5 human blastoids upon 50 nM ACBI1 treatment (ACBI1). Scale bar = 100 µm. **c** Box plot showing the change in SMARCA2 and SMARCA4 signal intensity in ACBI1-treated blastoids compared to DMSO, normalised by DAPI intensity, in day 5 blastoids. Data points are coloured by experiment. N=5 experiments. **d** Brightfield microscopic time-course imaging showing blastoid development in DMSO and ACBI1. Scale bar = 100 µm. **e** Plot showing the quantification of blastoid size over time upon ACBI1. Data points are shaped by experiment number. P-value is calculated by a multiple linear regression model on the log of the mean size. N=5 experiments. **f** Immunofluorescence images of day 2 and day 3 human blastoids showing trophectoderm marker GATA2, hypoblast marker GATA4, and epiblast marker SOX2. Scale bar = 100 µm. **g** Immunofluorescence images of day 5 human blastoids showing trophectoderm marker GATA2, epiblast marker SOX2, and hypoblast marker GATA4. Scale bar = 100 µm. **h** Violin plot showing the quantification of day 5 blastoid cell number. Data points are shaped by cell line and coloured by experiment. The lines represent the 25th, 50th, and 75th percentiles. N=7 experiments. **i** Violin plot showing the relative quantification of day 5 blastoid lineages based on the presence of marker genes and location of the cells. Data points are shaped by cell line and coloured by experiment. The lines represent the 25th, 50th, and 75th percentiles. N=8 experiments, n = 20423 (DMSO) and 26008 (ACBI1) cells. * = p-value < 0.05, ns indicates non-significance.

To investigate the effect of SWI/SNF perturbation on lineage acquisition, we performed immunofluorescence analysis on day 2 and day 3 human blastoids stained against EPI marker SOX2, TE marker GATA2, and HYPO marker GATA4. ACBI1 treatment increased the proportion of GATA2-positive cells from day 2 onwards **(Fig. 2f, Supplementary Fig. 4b).** At day 5, DMSO control blastoid size, total cell number, and lineage composition were comparable to those reported previously **(Fig. 2e, g-i)**^13^. We did not observe a difference in blastoid diameter and cell number between ACBI1-treated and DMSO-control blastoids at day 5 **(Fig. 2e, h)**. Nevertheless, these results suggest that degradation of SWI/SNF ATPases may alter lineage segregation during blastoid formation.

We therefore quantified lineage composition in day 5 blastoids by immunofluorescence imaging against SOX2, GATA2 and GATA4 using the machine learning function of Oxford Instruments’ Interactive Microscopy Image Analysis software (IMARIS). ACBI1-treated blastoids showed a marked reduction in SOX2-positive EPI cells and an increased proportion of GATA2-positive TE cells relative to the DMSO-treated controls **(Fig. 2g, i)**. Additionally, the number of GATA4-positive cells increased upon SWI/SNF perturbation **(Fig. 2g, i)**. Thus, degradation of SWI/SNF ATPases changes blastoid lineage composition, with a pronounced reduction of the SOX2-positive EPI cells and a relative increase of GATA2-positive TE cells.

We also examined non-cavitated structures to determine whether the lineage effect extended to aggregates that did not form blastoids. In the DMSO-treated controls, structures that failed to form blastoids were SOX2-positive and GATA2-negative **(Supplementary Fig. 4c).** By contrast, non-cavitated structures exposed to ACBI1 were predominantly GATA2-positive and SOX2-negative **(Supplementary Fig. 4c)**. These results are consistent with a shift from EPI to TE marker expression following SWI/SNF ATPase degradation, including in structures that fail to cavitate.

Together, these data show that SWI/SNF perturbation affects blastoid morphogenesis and lineage composition. Specifically, degradation of SWI/SNF ATPases compromises the formation or maintenance of the EPI lineage and is associated with an enhanced induction of the TE lineage.

To distinguish the contribution of SMARCA2 from the combined effect of SWI/SNF ATPase degradation, we generated blastoids treated with ACBI2, a PROTAC targeting only SMARCA2^56^. DMSO and ACBI1 were used as controls **(Supplementary Fig. 4d)**. Treatment with 5 nM ACBI2 was sufficient to induce more selective SMARCA2 degradation **(Supplementary Fig. 4e, f)**. Similar to ACBI1, ACBI2 treatment was associated with earlier cavitation, and ACBI2 treatment allowed for blastoid formation **(Supplementary Fig. 4g)**. We observed no difference in size upon ACBI2 treatment at day 3 and day 5 of blastoid formation **(Supplementary Fig. 4h)**. Overall, SMARCA2-specific degradation was associated with modest morphological changes.

We then analysed blastoid lineage composition at day 3 and day 5. SMARCA2-selective degradation was associated with a modest increase in GATA2-positive TE cells at day 3 and a reduction of SOX2-positive EPI cells at day 5 **(Supplementary Fig. 4i-k)**. These effects were less pronounced compared than those observed upon combined SWI/SNF ATPase degradation **(Supplementary Fig. 4i-k)**. Together, these results suggest that SMARCA2 degradation contributes to the lineage proportion phenotype, but that the more substantial effect of ACBI1 likely reflects the combined loss of both SWI/SNF ATPases. Thus, SMARCA2 contributes to the formation or maintenance of the EPI lineage during blastoid development.

### SWI/SNF ATPases degradation impairs blastocyst development and compromises ICM formation

We next investigated the role of SWI/SNF ATPases during early human embryonic development by treating human embryos cultured *ex vivo* with ACBI1. A total of 40, 3 dpf embryos at the 7-12 cell stage were cultured to 5 dpf in standard blastocyst medium, covering the period of the first lineage specification and blastocyst formation **(Fig. 3a)**. Embryos were treated with ACBI1 or DMSO as the control. Immunofluorescence analysis showed a significant reduction in SMARCA4, but not SMARCA2, protein levels **(Fig. 3b, c, and Supplementary Fig. 5a)**. The latter is consistent with the low abundance of SMARCA2 in 5 dpf human blastocysts **(Fig. 1c)**.

**Fig. 3:**
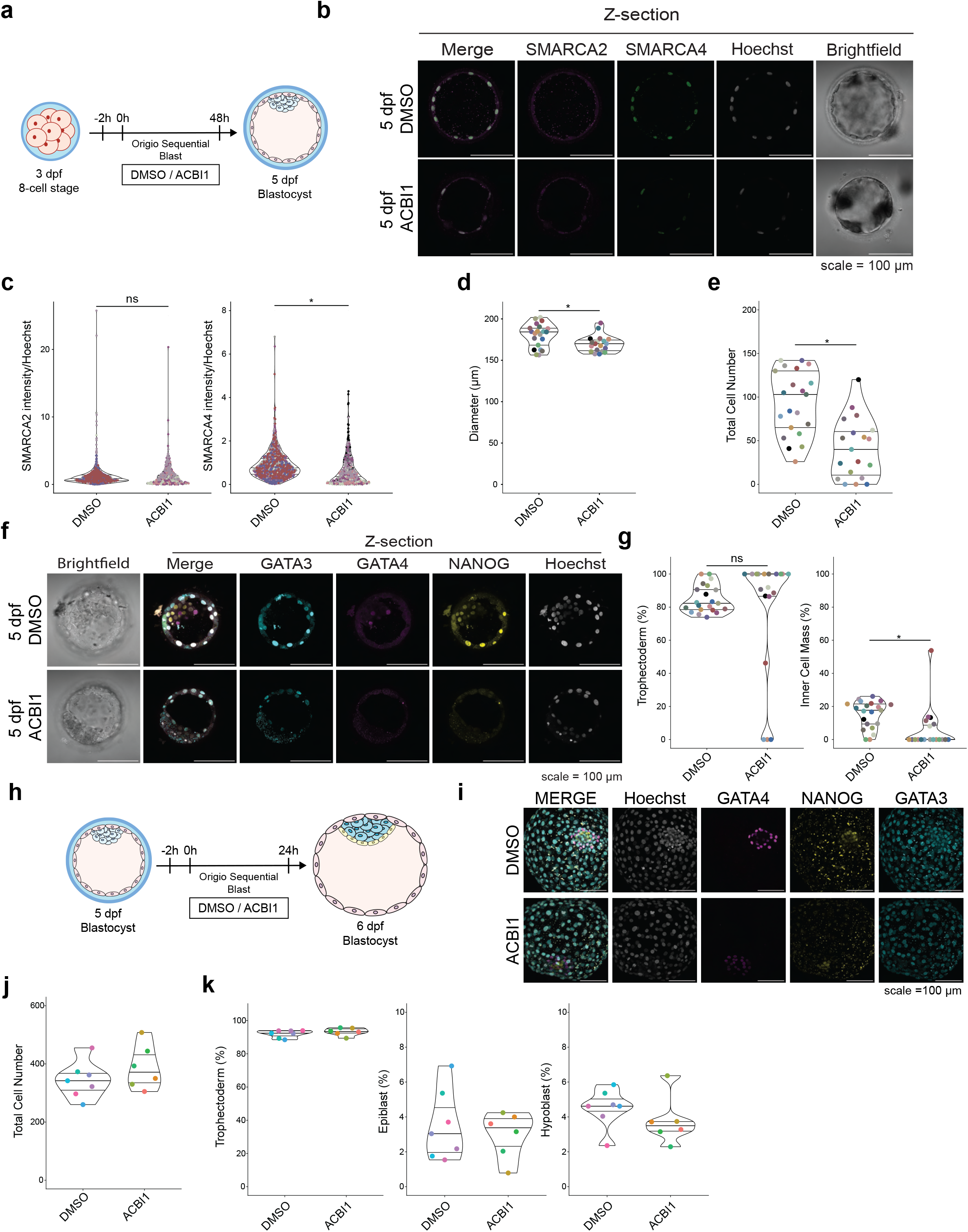
SWI/SNF ATPases safeguard blastocyst formation and cell fate acquisition. **a** Schematic outline of the experimental methodology for culturing human embryos from 3 to 5 days post-fertilisation (dpf) in blastocyst media containing either DMSO or 50 nM ACBI1. **b** Representative sections and z-projections of immunofluorescence-labelled embryos at 5 dpf for Hoechst (nuclear staining), SMARCA2 and SMARCA4. Scale bar = 50 μm. **c** Quantification of SMARCA2 and SMARCA4 normalised fluorescence intensity in embryonic cells in control and ACBI1-treated embryos. The lines represent the 25th, 50th, and 75th percentiles. N = 7 embryos per condition. **d** Violin plots showing the diameter of control and ACBI1-treated embryos at 5 dpf. The lines represent the 25th, 50th, and 75th percentiles. N = 21 (DMSO) and 19 (ACBI1) embryos. **e** Violin plots showing the quantification of total embryo cell number. The lines represent the 25th, 50th, and 75th percentiles. N = 21 (DMSO) and 19 (ACBI1) embryos. Quantification includes embryos not stained for lineage markers. **f** Representative sections and z-projections of immunofluorescence-labelled embryos at 5 dpf for Hoechst (nuclear staining), NANOG, GATA4 and GATA3. Scale bar = 50 μm. **g** Violin plots showing the quantification of trophectoderm (TE, left) and inner cell mass (ICM, right) lineage ratios. The lines represent the 25th, 50th, and 75th percentiles. N = 21 (DMSO) and 19 (ACBI1) embryos. Quantification includes embryos not stained for lineage markers. **H** Schematic outline of the experimental methodology for culturing human embryos from 5 to 6 dpf in blastocyst media containing either DMSO or 50 nM ACBI1. **I** Immunofluorescence-labelled embryos at 6 dpf for Hoechst (nuclear staining), NANOG, GATA4 and GATA3. Scale bar = 100 μm. **J** Violin plots showing the quantification of total embryo cell number. The lines represent the 25^th^, 50^th^, and 75^th^ percentiles. N = 7 (DMSO) and 6 (ACBI1) embryos. **K** Violin plots showing the quantification of trophectoderm (TE, left), epiblast (EPI, middle), and hypoblast (HYPO, right) lineage ratios. The lines represent the 25th, 50th, and 75th percentiles. N = 7 (DMSO) and 6 (ACBI1) embryos. * = p-value < 0.05, ns indicates non-significance.

ACBI1 treatment markedly impaired the quality of blastocyst development. Morphological assessment^57^ showed that 14 out of 21 DMSO-control blastocysts had an excellent or good ICM quality, compared to only one out of 19 ACBI1-treated blastocysts **(Supplementary Fig. 5b)**. The decrease in blastocyst quality was accompanied by a modest reduction in blastocyst diameter and a significant reduction in total cell number upon ACBI1 treatment **(Fig. 3d, e, Supplementary Fig. 5c)**. Overall, these results show that blastocyst development is affected by ACBI1 treatment.

The impairment of blastocyst quality was accompanied by a reduction in cell number across lineages **(Fig. 3f and Supplementary Fig. 5d, e)**. However, changes in lineage ratios appeared similar compared to ACBI1-treated human blastoids **(Fig. 3g and Fig. 2i)**. We next examined the effect of SWI/SNF perturbation on blastocyst lineage formation by performing immunofluorescence analsysis of TE marker GATA3, EPI marker NANOG, and HYPO marker GATA4 **(Fig. 3f and Supplementary Fig. 5d)**. We observed a significant reduction in GATA3-positive and NANOG-positive cells upon ACBI1 treatment **(Supplementary Fig. 5f)**. GATA4-positive cells were observed in 4 of 13 control embryos but were not detected following ACBI1 treatment **(Supplementary Fig. 5f)**, however, this result was not significant. A low abundance of HYPO cells is expected at the early blastocyst stage^3^. Immunofluorescence analysis of EPI marker SOX2, which shows some expression in early TE cells, revealed a significant reduction in SOX2 signal across the entire blastocyst following ACBI1 treatment **(Supplementary Fig. 5g, h)**. Altogether, these results suggest that SWI/SNF ATPases facilitate the specification of the TE and ICM lineages, especially safeguarding the formation of the ICM.

Notably, the effect of SWI/SNF ATPase degradation on NANOG-positive cells was more pronounced than on the number of ICM cells **(Fig. 3g and Supplementary Fig. 5e, f)**. For example, the only ACBI1-treated blastocyst with a morphologically well-defined ICM contained just four NANOG-positive cells among the 26 ICM cells. Additionally, we did not find GATA3-positive ICM cells, suggesting that degradation of SWI/SNF ATPases affects ICM lineage specification, but does not induce the TE fate within the ICM.

To investigate whether degradation of SWI/SNF ATPases affects the segregation of the EPI and HYPO lineages, we cultured 5 dpf embryos for 24 hours and treated them with DMSO or ACBI1 **(Fig. 3h)**. Degradation of the SWI/SNF ATPases was confirmed by immunofluorescence **(Supplementary Fig. 6a-c)**. Immunofluorescence analysis revealed no difference in cell number or cell lineage composition **(Fig. 3i-k and Supplementary Fig. 6d, e)**. Immunofluorescence intensities of NANOG and GATA4 were decreased in EPI and HYPO cells, respectively **(Supplementary Fig. 6f)**. However, these differences were not statistically significant after normalisation to the nuclear marker signal **(Supplementary Fig. 6g)**. Although 24-hour ACBI1 treatment might be too short to observe a phenotype, our results suggest that degradation of SWI/SNF ATPases does not induce or promote HYPO lineage formation in human blastocysts in this context. Whether EPI and HYPO maintenance is regulated by SWI/SNF ATPases remains unclear.

### Multiomic profiling reveals a misregulated TE state upon SWI/SNF ATPase degradation

Next, we aimed to dissect how SWI/SNF ATPases regulate early embryonic lineages. Given that multiomic profiling of human embryos is practically limiting, we performed 10X Genomics single-nucleus multiomic profiling of gene expression and chromatin accessibility on day 5 DMSO- and ACBI1-treated blastoids **(Fig. 4a)**.

**Fig. 4:**
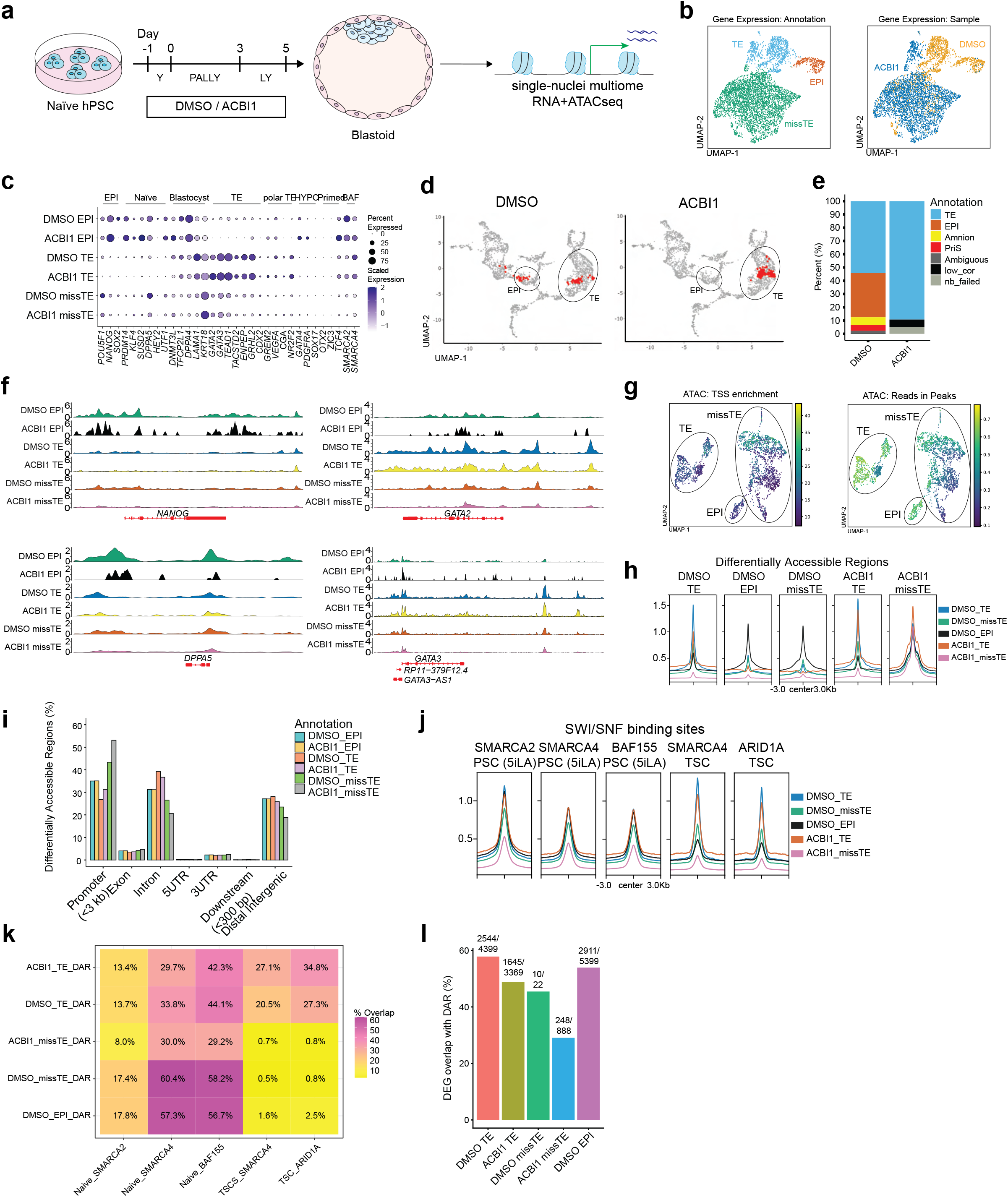
Multiomic profiling indicates that SWI/SNF ATPases establish blastoid cell identities through enhancer accessibility. **a** Schematic overview of the experimental set-up of 50 nM ACBI1 (ACBI1) during blastoid formation, followed by single-nucleus multiomic sequencing. **B** UMAP projections of the gene expression dataset coloured by cell annotation (left) and sample (right). **C** Dot plot showing expression of developmental marker genes per cell type and sample. Size represents the percentage of the group, and colour represents the scaled expression. **D** UMAP projections of gene counts integrated with an embryo reference dataset by^17^. Cells of this study are coloured in red. The trophectoderm (TE) and epiblast (EPI) clusters of the reference dataset are highlighted in red. **E** Stacked bar plot showing the number of cells annotated in each category by^17^. **F** ATAC Genomic tracks showing chromatin accessibility of epiblast markers *NANOG* and *DPPA5* and trophectoderm markers *GATA3* and *GATA2*. The genomic region is indicated in red. **G** UMAP coloured by transcription start site (TSS) enrichment score (left) and fraction of reads in peaks (right). **H** Profile plot showing chromatin accessibility over differentially accessible regions. **I** Bar plot showing the percentage of annotated genomic features of the differentially accessible regions per cell type and sample. **J** Profile plot showing chromatin accessibility over SWI/SNF binding sites identified in naïve hPSCs^42^ and TSCs^60^. **K** Heatmap showing the percentage of differentially accessible regions overlapping with SWI/SNF binding sites. **L** Bar plots displaying the percentage of differentially expressed genes overlapping with genes associated with differentially accessible regions.

Dimensionality reduction based on the transcriptome data identified three different clusters **(Fig. 4b and Supplementary Fig. 7a)**. One cluster expressed EPI markers such as *NANOG, DPPA4*, *DPPA5,* and *POU5F1,* which we therefore annotated as EPI **(Fig. 4b, c and Supplementary Fig. 7b, c)**. Another cluster expressed TE markers including *GATA3*, *TEAD1,* and *GATA2,* which we annotated as TE **(Fig. 4b, c, and Supplementary Fig. 7b, c)**. Expression analysis of early embryonic gene module genes identified by Meistermann et al.^49^, showed that DMSO-blastoid EPI cells were enriched for the EPI-specific POU5F1B and IFI16 gene modules, as well as the DUXA1 gene module **(Supplementary Fig. 7d)**. Blastoid TE cells showed a relative enrichment of TE-enriched GATA2 and NR2F2 gene modules **(Supplementary Fig. 7d)**. To benchmark human blastoids and cell type annotation, we integrated our blastoid dataset with a human embryo atlas^17^. This analysis revealed that DMSO-treated blastoid cells integrated mostly with the pre-implantation EPI and TE clusters of the human embryo reference **(Fig. 4d, e)**. In contrast, nearly all cells of ACBI1-treated blastoids integrated with the human embryo TE, and almost no EPI cells were detected **(Fig. 4d, e)**. These results are in line with our blastoid immunofluorescence analysis **(Fig. 2i and Fig. 3g)**. Altogether, our cell type annotation supports that our blastoids resemble the pre-implantation blastocyst stage^17^ and that the EPI is lost upon ACBI1-treatment.

In addition to the EPI and TE clusters, we identified a third cluster, which was enriched in the ACBI1-treated condition **(Fig. 4b, Supplementary Fig. 7c)**. This cluster displayed low expression of most EPI and TE marker genes compared to the EPI and TE clusters **(Fig. 4c and Supplementary Fig. 7b)**. Analysis of the Meistermann et al.^49^ gene module expression showed that these cells were enriched for the TE-associated DNMT3L and GATA2 gene modules **(Supplementary Fig. 7d)**. Furthermore, cells of this third cluster predominantly mapped to the reference human embryo TE, further supporting a TE-like transcriptome **(Fig. 4d, e)**. Given the integration with embryo TE, and expression of TE and EPI markers, we decided to refer to this transcriptionally distinct TE-like population as misspecified TE-like cells (missTE), while recognising that the data do not distinguish incomplete specification, altered maturation, or a stress-associated TE-like state **(Fig. 4b)**.

Next we aimed to further characterise missTE cells. We examined immunofluorescence images of GATA3 and TEAD1 in blastoids collected from the same batch as the multiomic dataset. Our analysis revealed that both TEAD1 and GATA3 remained detectable at the protein level irrespective of the blastoid phenotype and condition **(Supplementary Fig. 7e)**. Gene set expression analysis revealed no clear misregulation of apoptosis, senescence, or cell cycle genes in missTE cells **(Supplementary Fig. 7f)**. Differential gene expression analysis showed that missTE cells were enriched for genes related to oxidative phosphorylation and translation and down-regulated genes related to DNA damage, transcription, and cell signalling **(Supplementary Table 1)**. In addition to ribosomal and oxidative phosphorylation proteins, missTE cells displayed a relatively high expression of housekeeping genes^58^ **(Supplementary Fig. 7g)**. In contrast, ACBI1-TE cells upregulated genes related to Nuclear transport, RNA biosynthesis regulation, and chromatin remodelling **(Supplementary Table 1)**. Together, these analyses suggest that missTE cells represent a transcriptionally distinct TE-like population, although the exact identity and origin remain unclear.

Overall, our blastoid single-nucleus transcriptome analysis shows that ACBI1 treatment is associated with a loss of the EPI fate and a gain of a TE-like identity. Consistent with the blastoid immunofluorescence data, this indicates that SWI/SNF ATPase degradation is associated with a blastoid lineage composition depleted in EPI cells and favouring a TE-like state.

### SWI/SNF ATPase degradation is associated with impaired accessibility at enhancer regions

We then investigated the impact of SWI/SNF ATPase degradation on chromatin accessibility in human blastoids, by performing pseudobulk analysis on the single-nucleus ATAC data based on their respective sample and cell type annotations **(Supplementary Fig. 7h)**. DMSO-EPI cells showed increased chromatin accessibility over EPI genes such as *NANOG* and *DPPA5,* while DMSO-TE cells showed increased accessibility over TE genes such as *GATA2* and *GATA3,* as expected **(Fig. 4f)**. ACBI1-TE cells showed a similar accessibility pattern over TE genes compared to DMSO-TE cells **(Fig. 4f)**. MissTE cells, specifically ACBI1-missTE cells, showed a markedly reduced chromatin accessibility at distal regions, while promoter peaks were maintained at these genes **(Fig. 4f)**. Furthermore, ACBI1-missTE cells exhibited increased transcription start site (TSS) enrichment with fewer reads in peaks, consistent with reduced accessibility at distal regulatory regions upon the loss of SWI/SNF ATPases **(Fig. 4g)**.

Next, we identified differentially accessible regions (DARs) for each cluster relative to all other clusters, using pyCisTopic^59^. DMSO-TE and EPI clusters displayed lineage-specific accessibility **(Fig. 4h)**. Interestingly, DMSO-TE cells had increased accessibility over ACBI1-TE DARs **(Fig. 4h)**. Additionally, DMSO-missTE DARs were mostly accessible in DMSO-EPI cells, while ACBI1-missTE DARs were accessible in all other conditions, indicating an altered, less lineage-restricted chromatin accessibility profile in ACBI1-missTE cells, while DMSO-missTE cells maintain a slight EPI-like profile **(Fig. 4f, h)**. Overall, this analysis suggests that ACBI1 treatment is associated with impaired chromatin remodelling.

Moreover, ACBI1-blastoid cells showed a relative increase in promoter accessibility and a reduction in accessibility at distal intergenic regions, in both peaks and DARs compared to DMSO-blastoid cells **(Fig. 4f, i and Supplementary Fig. 7i-k)**. This trend was also observed when comparing the ACBI1-missTE to the DMSO-missTE cells, or ACBI1-TE to DMSO-TE cells **(Fig. 4i and Supplementary Fig. 7k).** Consistent with these findings, topic modelling^59^ independently revealed similar shifts in chromatin accessibility following SWI/SNF ATPase degradation **(Supplementary Fig. 7l, m and Supplementary Table 2).** Taken together, these results suggest that SWI/SNF ATPase degradation leads to a loss of chromatin accessibility at distal regions, while promoter accessibility was relatively preserved.

To determine whether the chromatin accessibility changes induced by SWI/SNF ATPase degradation occurred specifically at regions directly occupied by SWI/SNF complexes, we next analysed chromatin accessibility over SWI/SNF binding sites identified in published ChIP-seq and CUT&RUN datasets^42,60^. Our analysis revealed that SWI/SNF-bound regions in naïve hPSCs were equally accessible in TE cells, while SWI/SNF binding sites in hTSCs were specific to only TE cells **(Fig. 4j)**. Although ACBI1-TE cells showed a similar pattern to DMSO-TE cells, ACBI1-missTE cells showed a strong reduction in SWI/SNF binding-site accessibility **(Fig. 4j, k)**. Altogether, these results suggest that the missTE state was associated with reduced accessibility at SWI/SNF-bound regions.

Next, we compared differential gene expression with differential accessibility. ACBI1-treated cells showed reduced overlap between differential expression and gene accessibility **(Fig. 4l)**. Overall, these results suggest that the degradation of SWI/SNF ATPases was associated with a reduced correlation between chromatin accessibility and gene expression.

### Integrative analysis reveals SWI/SNF ATPase degradation is correlated with impaired accessibility at predicted regulatory elements

Next, we aimed to investigate the impact of SWI/SNF ATPases on the gene regulatory programs in blastoids. We used SCENIC+ to infer TFs, candidate target enhancers, candidate target genes, and enhancer-driven regulons (eRegulons) **(Fig. 5a and Supplementary Fig. 8a)**. The combination of these eRegulons in a cell forms enhancer-driven gene regulatory networks (eGRNs)^22^. Analysis of the DMSO-blastoids dataset alone (indicated by the asterisk in **Fig. 5b-d**) revealed 105 eRegulons, of which 102 were identified in at least 5/10 SCENIC+ iterations (see Methods) **(Fig. 5b-d, Supplementary Table 3)**. Analysis of the combined dataset revealed 355 eRegulons, of which 28 were identified in at least 5/10 iterations **(Supplementary Fig. 8b, c, Supplementary Table 3)**. In total, we identified 50,550 enhancer regions, associated with 11,643 target genes **(Supplementary Table 4)**. This dataset contains regulons that were previously identified in human embryos using SCENIC^23^, as well as novel regulons associated with distinct blastoid lineages **(Fig. 5a, b and Supplementary Fig. 8b, c and Supplementary Tables 3, 4)**.

**Fig. 5:**
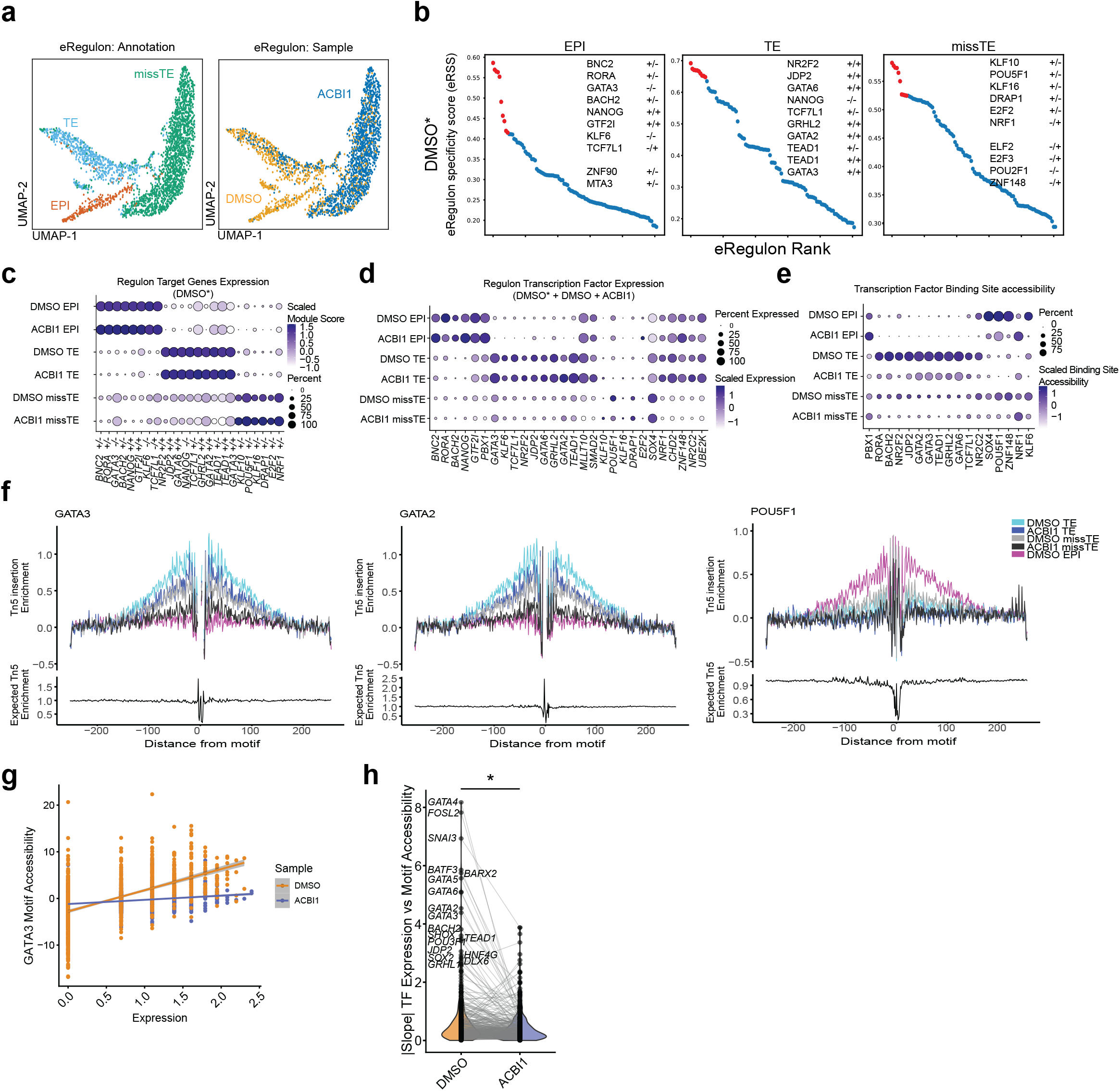
Gene regulatory network analysis reveals SWI/SNF ATPases facilitate transcription factor motif accessibility. **a** UMAP projections of the eRegulon dataset coloured by cell annotation (left) and sample (right). **b** Plot showing eRegulon specificity score (eRSS) of eRegulons ranked per cluster. The top 10 regulons are shown in red. * = eRegulons predicted from analysis of the DMSO condition alone. **c** Dot plot showing the expression of eRegulon target genes per cell type and sample. Size represents the percentage of the group, and colour represents the scaled module score. * = eRegulons predicted from analysis of the DMSO condition alone. **d** Dot plot showing expression of eRegulon transcription factors per annotation and sample. Size represents the percentage of the group, and colour represents the scaled expression. **e** Dot plot showing the accessibility score of TF motifs (JASPAR2020) of SCENIC+-predicted transcription factors per annotation and cell type. Size represents the percentage of the group, and colour represents the scaled motif-accessibility score. **f** Transcription factor footprint analysis of transcription factors GATA2, GATA3, and POU5F1. **g** Scatterplot showing the relationship between *GATA3* expression (x-axis) and motif accessibility (y-axis). **h** Violin plot showing the absolute value of the slope between expression and motif accessibility per transcription factor. N = 300 genes. * = p-value < 0.05.

Predicted EPI-associated eRegulons included NANOG^23,61^ and PBX1^23^, together with BNC2, RORA, BACH2, and GTF2I **(Fig. 5b, Supplementary Fig. 8b)**, while eRegulons associated with the TE fate included NR2F2^23^, JDP2, GATA2^23^, GATA3^23^, TEAD1^23^, as well as GRHL2, GATA6, TCF7L1, NR2C2, SMAD2, and MLTT10 **(Fig. 5b, and Supplementary Fig. 8b)**. The inferred network predicted negative regulatory relationships between GATA3, TCF7L1, TEAD1, and KLF6 and EPI-associated regulatory programs while NANOG-associated regulon activity was predicted to oppose TE-associated regulatory programs **(Fig. 5b and Supplementary Fig. 8b, d)**. MissTE cells were associated with the eRegulon activity of POU5F1, SOX4, and GATA3, but are also marked by the lack of the repressive activity from TEAD1, NR2C2, UBE2K, ZNF148, NRF1, and CHD2 **(Fig. 5b, c and Supplementary Fig. 8b, c)**. Altogether, our analysis predicts candidate TFs and regulatory programs underlying human blastoid lineage identity.

Next, we generated a gene regulatory network (GRN) from the predicted eRegulons, suggesting *NANOG* as a key regulator of the EPI GRN. We predicted an interconnected TE GRN where TEAD1, GATA3, and TCF7L1 inhibit the drivers of EPI identity **(Supplementary Fig. 8d)**. The inferred network predicted NANOG, PBX1, and ZNF148, as candidate positive regulators of *SMARCA2*, and ZNF148 and SMAD2 as positive regulators of *SMARCA4* **(Supplementary Fig. 8d)**. MissTE cells were characterised by reduced expression of multiple TFs represented in the inferred network **(Fig. 5d)**, suggesting the SWI/SNF complex plays a role in enhancer-driven expression of developmental TFs.

Although some difference was observed in the eRegulon specificity score (RSS) per eRegulon between the DMSO and ACBI1 lineages **(Fig. 5b and Supplementary Fig. 8b)**, little difference could be observed in the expression of their target genes **(Fig. 5c and Supplementary Fig. 8c)**. Quantification of eRegulon-TF motifs accessibility revealed that even though ACBI1-TE cells expressed TE-associated TFs similarly to DMSO-TE cells **(Fig. 5d)**, ACBI1-TE cells showed a decrease in TF motif accessibility **(Fig. 5e)**. This suggests that TE cells have defective chromatin remodelling at motif sites targeted by lineage-associated TFs upon the degradation of SWI/SNF ATPases.

To further investigate the effect of SWI/SNF ATPase degradation on TF target site accessibility, we performed TF-footprinting analysis^62,63^. GATA2 and GATA3 footprint was highest in DMSO-TE cells, while the POU5F1 footprint was highest in DMSO-EPI cells **(Fig. 5f)**. Interestingly, the GATA2 and GATA3 footprint of ACBI1-TE cells closely resembled that of DMSO-missTE cells, while the footprint of ACBI1-missTE cells were similar to that of DMSO-EPI cells **(Fig. 5f)**. Accessibility analysis of SCENIC+-predicted TF-targeted enhancer regions showed that predicted NANOG-targeted enhancers were only accessible in DMSO-EPI cells. Furthermore, enhancer regions targeted by GATA2 and GATA3 in ACBI1-TE cells showed lower accessibility compared to DMSO-TE cells **(Supplementary Fig. 8e)**. However, accessibility of GATA3 binding sites in human TSCs^64^ did not show this pattern **(Supplementary Fig. 8f)**. Interestingly, even though global promoter accessibility was generally maintained, accessibility of SCENIC+-predicted GATA3 target sites was also decreased within promoter regions upon ACBI1 treatment **(Supplementary Fig. 8g)**. In DMSO-blastoids, upregulation of GATA3 expression was correlated with an increase in GATA3 motif accessibility **(Fig. 5g)**. This correlation was lost upon ACBI1 treatment. Further analysis revealed a general decrease in correlation between TF expression and target motif accessibility upon ACBI1-treatment **(Fig. 5g, h and Supplementary Fig. 8h, i)**. Altogether, our results suggest that SWI/SNF ATPases facilitate the accessibility of regulatory regions targeted by lineage-associated TFs in human blastoids.

### SWI/SNF ATPases regulate the specification and maintenance of human blastoid lineages

To dissect the role of SWI/SNF ATPases in lineage dynamics, we utilised the acute nature of the PROTAC system by treating blastoids for different time-points and durations with ACBI1 **(Fig. 6a)**. To assess whether TE induction resulted from a loss of pluripotency, we treated blastoids with ACBI1 during aggregation only (ACBI1 -1:0). To differentiate the role of SWI/SNF ATPases in lineage specification and maintenance, we treated blastoids until day 3 (ACBI1 -1:3), or from day 3 (ACBI1 3:5) of blastoid formation, respectively. SWI/SNF ATPase degradation was validated by immunofluorescence at day 3 and day 5 of blastoid formation **(Supplementary Fig. 9a, b)**, although SMARCA2 and SMARCA4 levels did not completely recover after 2 days in ACBI1 -1:3. All conditions supported blastoid formation, and we found no significant difference in aggregate size at either day 3 or day 5 **(Supplementary Fig. 9c, d)**.

**Fig. 6:**
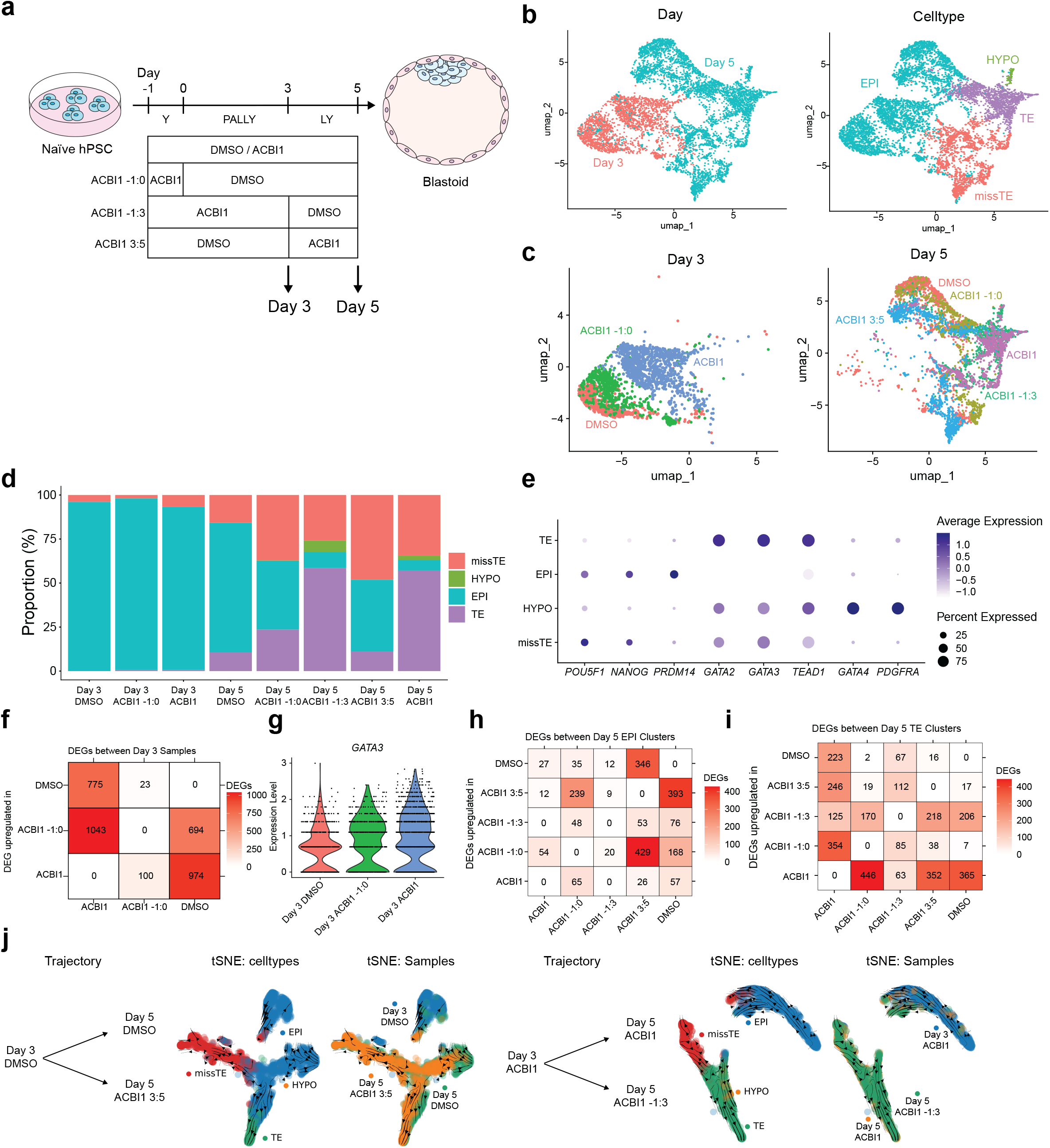
Single-nucleus transcriptome sequencing reveals SWI/SNF ATPases affect cell lineage establishment and trajectories. **a** Schematic overview of the experimental set-up of 50 nM ACBI1 (ACBI1) during blastoid formation, followed by single-nucleus transcriptome sequencing by PIPseq. **B** UMAP projections coloured by day (left) and cell type (right). **C** UMAP projections coloured by sample of day 3 (left) and day 5 (right) blastoids. **D** Stacked bar plot showing cell type proportion per sample. **E** Dot plot showing expression of developmental marker genes per cell type. Size represents the percentage of the group, and colour represents the scaled expression. **F** Heatmap showing the number of upregulated genes between day 3 samples. The number shows the number of upregulated genes in the row versus the column. **G** Violin plot showing normalised GATA3 expression in day 3 samples. **H** Heatmap showing the number of upregulated genes between day 5 EPI clusters. The number shows the number of upregulated genes in the row versus the column. **I** Heatmap showing the number of upregulated genes between day 5 TE clusters. The number shows the number of upregulated genes in the row versus the column. **J** CellRank2 pseudotime analysis showing cell trajectories between day 3 and day 5.

Blastoids treated with ACBI1 during the aggregation stage displayed enhanced cavitation at day 3 **(Supplementary Fig. 9c)**. Immunofluorescence analysis of GATA2, SOX2, and GATA4 at this stage revealed that blastoids treated with ACBI1 during aggregation displayed an intermediate induction of the TE fate at day 3 **(Supplementary Fig. 9e, f)**. These results suggest that enhanced TE induction upon ACBI1 treatment is partly the result of a changed pluripotent state at the start of blastoid induction.

Immunofluorescence analysis of day 5 blastoids showed that ACBI1 treatment during aggregation and during the last 2 days of blastoid formation did not lead to altered EPI formation **(Supplementary Fig. 9f)**, reminiscent of our results in blastocysts **(Fig. 3h-k)**. Day 5 blastoids treated with ACBI1 until day 3 displayed perturbed EPI formation, similar to blastoids that received continuous treatment **(Supplementary Fig. 9f)**. Altogether, our results suggest that SWI/SNF ATPases regulate lineage specification in blastoids.

To investigate lineage dynamics at the single-cell level, we performed single-nucleus RNA-seq on day 3 and day 5 blastoids treated with ACBI1 for different durations. We obtained sequencing data from 7246 nuclei, with a median of 811 nuclei per sample. Our analysis identified EPI, TE, missTE, and HYPO populations, expressing their respective marker genes. Furthermore, our time-course dataset could be integrated with our multiome dataset and the human embryo reference atlas^17^ **(Fig. 6b-e and Supplementary Fig. 10a-c)**.

All ACBI1-treated blastoid conditions showed an enrichment in missTE cells, although the proportion of missTE over TE cells was decreased compared to the multiome dataset **(Fig. 6d)**. In line with the immunofluorescence analysis, ACBI1 and ACBI1 -1:3 blastoids showed a decrease in the number of EPI cells **(Fig. 6d)**. Most day 3 cells showed an EPI-like identity in all conditions **(Fig. 6d and Supplementary Fig. 10d and Supplementary Table 5)**. However, both ACBI1-treated day 3 conditions showed a large number of DEGs compared to the DMSO condition **(Fig. 6f)**. Furthermore, *GATA3* expression was increased in both ACBI1-treated conditions at day 3 **(Fig. 6g and Supplementary Fig. 10e)**, consistent with our immunofluorescence analysis **(Supplementary Fig. 9f and Supplementary Fig. 10f)**. Differential Gene Expression analysis between the different EPI and TE clusters revealed that ACBI1-treatment between day 3 and day 5 lead to transcriptional misregulation in EPI cells **(Fig. 6h, Supplementary Table 5)**, while the misregulation of the TE lineage was more apparent in cells treated with ACBI1 before compared to after day 3 **(Fig. 6i, Supplementary Table 5)**. Altogether, these data support that SWI/SNF ATPases establish and maintain the EPI while also regulating TE establishment in blastoids.

To assess the influence of SWI/SNF ATPase perturbation on developmental trajectories, we performed pseudotime trajectory analysis using CellRank2^65–67^. This analysis placed missTE cells on a trajectory distinct from the principal EPI and TE trajectories, rather than an apparent intermediate position between these states **(Fig. 6j and Supplementary Fig. 10h)**. This may suggest that the missTE identity is an alternative cellular state that is enriched upon the loss of SWI/SNF ATPases **(Fig. 6d and Supplementary Fig. 7c)**.

Interestingly, day 3 blastoids treated with ACBI1 for three days showed an increased fate probability for the TE fate compared the DMSO and ACBI1 -1:0 blastoids **(Supplementary Fig. 10h)**. Furthermore, trajectory analysis suggested missTE cells in ACBI1 -1:0 and ACBI1 3:5 blastoids branched from an EPI-like state, while missTE cells in continuous ACBI1 and ACBI1 -1:3 blastoids were predominantly connected to a TE-like state **(Fig. 6j and Supplementary Fig. 10h)**. These results suggest that missTE cells have distinct precursor states across ACBI1 treatment windows.

Altogether, dynamic degradation of SWI/SNF ATPases during blastoid formation suggested a stage-dependent role of SWI/SNF ATPases in blastoid lineage dynamics. Early SWI/SNF ATPase degradation was associated with enhanced specification of the TE fate, transcriptional misregulation of TE cells, enrichment of the missTE state, and loss of the EPI. Late SWI/SNF ATPase degradation was associated with transcriptional misregulation within EPI cells. These results suggest that the SWI/SNF complex regulates lineage specification and maintenance during blastoid formation.

In conclusion, degradation of SWI/SNF ATPases was associated with a decrease in TF motif accessibility and impairments in the establishment and maintenance of the TE and EPI fates, suggesting that the SWI/SNF complex regulates the development of lineages during human pre-implantation development **(Figure 7)**.

**Fig. 7:**
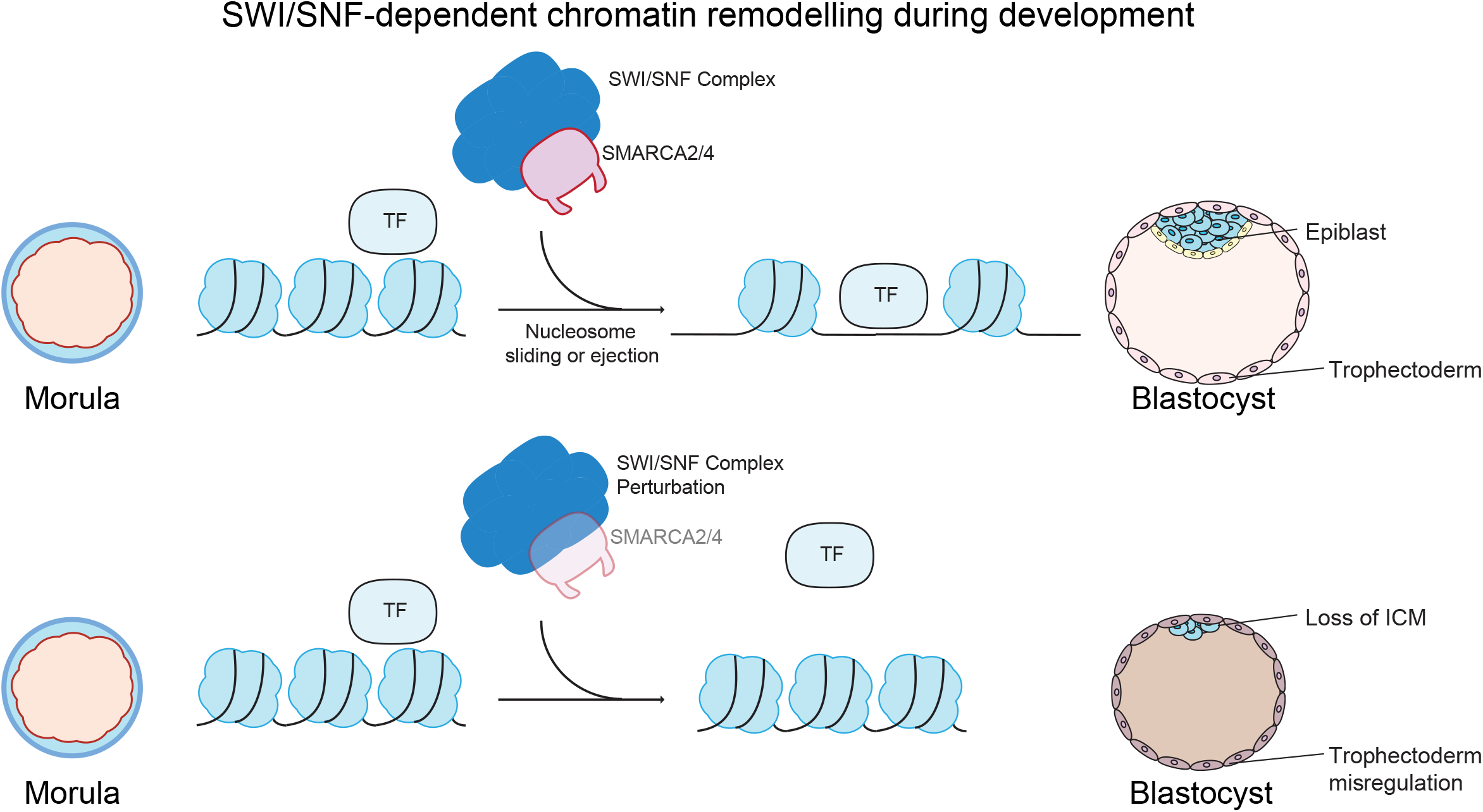
Schematic representation of SWI/SNF regulation during human pre-implantation development. During pre-implantation development, the human embryo specifies the first cell lineages, segregating the embryonic epiblast from the extraembryonic trophectoderm. During this event, the SWI/SNF complex interacts with transcription factors to facilitate chromatin accessibility. Upon the removal of SWI/SNF ATPases, the epiblast identity will not be maintained, and the establishment of the trophectoderm fate by the trophectoderm regulatory program is impaired.

## Discussion

The establishment of cell fates during early development is accompanied by dramatic changes in the chromatin landscape^27^. These changes are regulated by the SWI/SNF complex, which interacts with TFs at cis-regulatory elements to ensure the establishment of gene regulatory programs^25–28^. However, the role of the SWI/SNF complex in early pre-implantation human embryogenesis had not yet been revealed. In this study, we investigated the role of SWI/SNF ATPases in early human embryonic development in three model systems, both *ex vivo* and *in vitro*. Our analysis shows that SWI/SNF ATPases have distinct developmental expression patterns. Moreover, the SWI/SNF complex facilitates TE differentiation while also safeguarding ICM and EPI formation in adherent culture, blastoids, and human blastocysts. Using single-nucleus multiomic and transcriptomic profiling, our data revealed that upon the removal of SWI/SNF ATPases, cells displayed decreased TF motif accessibility, correlating with improper TE establishment. Altogether, our data suggest that SWI/SNF ATPases facilitate the establishment of cell identities in human blastoid and pre-implantation development, and regulate cis-regulatory element accessibility.

Although the role of the SWI/SNF complex in murine pre-implantation development is well established^34–41^, it was previously unexplored in human pre-implantation development. Murine embryos require *Smarca4*, but not *Smarca2,* to develop past the implantation stage^34,38^ and perturbation of *Smarca4* or other core SWI/SNF proteins leads to upregulation of pluripotency markers in TE cells of murine blastocysts, which displayed no morphological differences^34–40^. Additionally, *Smarca4*-deficient blastomeres preferentially contribute to the ICM^39^. In our study, we found that the expression dynamics of *SWI/SNF ATPases* in human blastocysts resemble the expression patterns described in murine blastocysts, at least with regard to lineage specificity^68^. However, in contrast to mice, where blastocyst morphology is unaffected, we found that human blastocyst development, and specifically ICM formation, is impacted by SWI/SNF ATPase perturbation. Furthermore, we did not observe an upregulation of pluripotency factors in TE cells of adherent cells, blastoids or blastocysts upon SWI/SNF degradation. These results show that regulation of development by the SWI/SNF complex is not entirely conserved between mice and humans.

Differences in the regulation of naïve pluripotency by the SWI/SNF complex between mice and humans have previously been described^42^. While murine naïve pluripotency seems almost entirely dependent on *SMARCA4*^36,37^, human naïve pluripotency is regulated by both SWI/SNF ATPases, and combined, but not individual, depletion of both SWI/SNF ATPases results in loss of human naïve pluripotency^42^. Here, we confirmed that the SWI/SNF complex is required for maintenance of human naïve pluripotency. However, in contrast to *SMARCA2* knockout combined with SMARCA4 knockdown^42^, we did not observe a collapse of colony morphology, but only slightly smaller colonies. This might be explained by differences in culture conditions or due to a different method of SWI/SNF perturbation (genetic versus protein depletion). In addition to regulating the maintenance of naïve pluripotency, we found that the formation of the ICM and EPI is impaired upon loss of SWI/SNF ATPases in both human blastoids and blastocysts. By showing that the SWI/SNF complex safeguards the formation of the ICM and the EPI compartments in blastocysts and blastoids, we extend our previous knowledge on the role of the SWI/SNF complex in the maintenance of naïve pluripotency and human development.

We extended our investigation on the role of SWI/SNF ATPases in TE specification across three complementary systems: adherent cell differentiation, blastoid formation and human blastocyst development. Although SWI/SNF perturbation impaired TE establishment across all three systems, we also observed system-specific phenotypic differences. Such differences between adherent and blastoid differentiation have been reported previously^69^, and may reflect variations in the cellular, signaling and culture contexts in which differentiation occurs^6,13^. Additionally, pre-implantation human embryo development requires little to no exogenous signalling modulation to specify the embryonic lineages^70^. The absence of exogenous signalling-modulating cues could explain the lack of extraembryonic lineage induction within ICM cells, while blastoids enhance TE and HYPO formation upon ACBI1-treatment. This suggests that SWI/SNF ATPases regulate the conversion of cell fates and the establishment of cell identities, rather than inducing differentiation. Altogether, even though phenotypic differences between these systems exist, we found that the SWI/SNF complex facilitates the establishment of the TE fate.

Previous studies show that the SWI/SNF complex is necessary to maintain the accessibility of most enhancers, while promoter accessibility is restored within four hours of SWI/SNF inhibition in murine embryonic stem cells and mammary gland cells and that the epigenetic landscape and cis-regulatory elements affect SWI/SNF-dependence^31,33^. In our study, we observed impaired enhancer accessibility upon SWI/SNF perturbation, while accessibility at promoter regions was maintained. Furthermore, we observed impaired TF motif accessibility upon the removal of SWI/SNF ATPases. How the TE program is activated during SWI/SNF ATPase degradation remains to be investigated. However, we hypothesise that TE-driving TFs, such as GATA2, GATA3, TEAD1, GRHL2, JDP2, and NR2F2, may depend on SWI/SNF ATPases for effective transcriptional activity by maintaining enhancer and motif accessibility required for the establishment of the TE fate.

In line with the loss of enhancer accessibility and promoter accessibility upon acute SWI/SNF perturbation^31,33^, we found that promoter accessibility was maintained, even after 6 days of SWI/SNF degradation, compared to the 4-30 hours of inhibition in previous studies^31,33^. Additionally, Basurto-Cayuela and colleagues described how housekeeping genes were SWI/SNF-independent^33,58^, which could be explained by their chromatin profile and lack of enhancer regulation^31,33,71^. We found that cells that displayed impaired enhancer accessibility retained expression of housekeeping, ribosomal protein, and oxidative phosphorylation genes. Altogether, our data suggest that the SWI/SNF complex specifically regulates accessibility of enhancers and TF motifs to specify and maintain cell identity, but not for the expression of housekeeping genes.

In conclusion, our study provides evidence for the essential role of the SWI/SNF complex in safeguarding human preimplantation development, supported by *ex vivo* human blastocyst experiments. Together, our findings underline the importance of chromatin processes in early human development, provide a valuable framework for understanding how chromatin remodelling contributes to cell fate acquisition in human embryogenesis, and provide further insight into how stem cell embryo models recapitulate and differ from human embryo development.

## Future Perspective

How cells still differentiate towards the TE fate and open or maintain chromatin accessibility upon SWI/SNF perturbation remains unclear. It is possible that chromatin or other epigenetic mechanisms compensate for or react to the loss of SWI/SNF ATPases and therefore allow for differentiation into the TE fate. Furthermore, we showed that TF motif accessibility is decreased upon depletion of SWI/SNF ATPases. However, whether enhancer activity is also decreased upon ACBI1 treatment remains to be systematically investigated. Further investigation will be needed to shed light on the molecular events and dynamics underlying SWI/SNF-mediated cell fate establishment, and its role in post-implantation development.

## Limitations

In our study, we used naïve hPSCs to nTE conversion, blastoids, and early human embryos to investigate the role of SWI/SNF ATPases in human embryogenesis. While blastoids model the blastocyst stage, we recognise that blastoids used in this study represent a later blastocyst stage than the blastocysts used in this study. In our study, we compared blastoid formation to the development of 3 dpf embryos to 5 dpf embryos. Caution should be used when comparing blastoid formation with early blastocyst development, as blastoid development is the differentiation of the three founding lineages from EPI-like cells *in vitro*, while in human embryos, ICM and the TE cells are specified from pre-lineage morula cells. While valuable, stem cell-based embryo models are close but not exact replicas of embryos. Although this study does not include a multiomic and transcriptomic dataset for human blastocysts, we believe that our blastoid datasets provide valuable insights into the role of the SWI/SNF complex in human development.

We acknowledge our use of different markers to assess the TE and EPI identities in human blastocysts and blastoids, mainly due to practical considerations. However, given the similarity between GATA2 and GATA3, and SOX2 and NANOG at the RNA^17^ and protein^49,72^ level in the blastocyst stage, we do not foresee problems with our interpretation of the data.

To perturb the SWI/SNF complex, we used the PROTAC ACBI1^54^. ACBI1 allows for a fast and inducible degradation of SWI/SNF ATPases. However, ACBI1 treatment does not remove all proteins and can therefore be compared to a knockdown, rather than a full knockout. By degradation of the SWI/SNF ATPases, the SWI/SNF complex is perturbed; however, not through catalytic inhibition alone. With our results, we are not able to entirely distinguish between the specific role of SMARCA2 and SMARCA4, or between catalytic and non-catalytic effects, including the recruitment of other chromatin factors. Additionally, we are unable to distinguish acute from indirect effects of SWI/SNF perturbation on cell identity, gene expression, or chromatin accessibility^31,33,73^. Furthermore, even though our data show a correlation between SWI/SNF perturbation, loss of TF motif accessibility, and transcriptomic and developmental misregulation, we have not directly assessed SWI/SNF binding or the effect of SWI/SNF perturbation on TF activity and enhancer activity in this study.

## Materials and Methods

### Cell culture

#### Feeder-free culture of human naïve pluripotent stem cells

Naïve hPSCs (iPSC EPITHELIAL-1 IPSC0028, Sigma-Aldrich (SIGi001-A), converted to the naïve state by KLF4 overexpression^74^, WA09, WiCell (H9), converted to the naïve state by HDAC inhibition^75^, and CAMe001-A (HNES1)) were cultured at 37°C in hypoxic conditions (5% O_2_, 5% CO_2_) with filter sterilised N2B27 medium consisting of 48% DMEM/F-12 (Gibco, 31330-028), 48% Neurobasal (Gibco, 21103-049), 0.5X Glutamax (Gibco, 35050-038), 0.5% N2-supplement (Gibco, 17502-048), 1% B27-supplement (Gibco, 17504-044), 1X Non-essential amino acids (NEAA) (Gibco, 11140-035), 1% Penicillin-Streptomycin (Gibco, 15140-122) and 0.05 mM *β*-mercaptoethanol (Gibco, 31350-010), freshly supplemented with 1 µM PD0325901 (Axon Medchem, 1408), 2 μM XAV939 (Sigma, X3004), 2 μM Gö6983 (Tocris, 2285), and 10 ng/mL human LIF (PeproTech, 300-05-5UG) to make PXGL medium^50^, which was changed daily.

Naïve hPSCs were passaged every three days by single-cell dissociation using Accutase (Sigma, A6964-500ML) (5 minutes at 37°C). Plates were prepared the day before passaging and incubated at 37°C overnight with FBS-coating media consisting of 87% DMEM/F-12, 10% FBS (Gibco, 10270), 1% Penicillin-Streptomycin and 0.1 mM *β*-mercaptoethanol^76^. On the day of passaging, 10 µM of Y-27632 (Tocris, 1254) was added to the PXGL medium.

When naïve hPSCs were cultured on mouse embryonic fibroblasts (MEFs) feeders as previously described^43^, they were converted to feeder-free culture by plating them on FBS-coated plates, as described above. Cells were cultured feeder-free for a few passages before they were used for an experiment.

Cells were regularly tested for mycoplasma contamination. The cell line stocks used were short-tandem repeat (STR)-authenticated and karyotyped at the University Hospital of Leuven (UZ Leuven).

#### Naïve-to-trophectoderm conversion

The conversion of feeder-free naïve hPSCs to TE was performed using a protocol adapted from Io et al. (2021)^6^. Naïve hPSCs were cultured in PXGL prior to conversion. On day 0, cell culture plates (Corning,3516) were coated with 0,15 µg/cm^2^ iMatrix Silk Laminin 511 (AMSBIO, AMS.892 021) in 1X PBS (Gibco, 10010023) and incubated for at least 30 minutes at 37°C. Around 4 × 10^4^ cells/cm² were plated and cultured in N2B27 media supplemented with 2 µM PD0325901, 2 µM A83-01, and 10 ng/ml BMP4 (Biotechne, 314-BP-010). From day 1 onward, cells were cultured in N2B27 supplemented with 2 µM PD0325901, 2 µM A83-01, and 1 µg/ml JAKi1 (Merck, 420099-500UG). The media was changed on day 2.

#### N2B27 culture

Naïve hPSCs were cultured in PXGL prior to culture in N2B27. FBS-coated plates were prepared the day before passaging and incubated at 37°C overnight. On day 0, cells were dissociated into single cells using Accutase. Cells were counted, and around 1.5-2.5 × 10^4^ cells/cm² were plated and cultured in N2B27 supplemented with 10µM of Y-27632 (Tocris, 1254) to form N2B27Y. From day 1 onward, cells were cultured in N2B27. Cells were collected on day 3.

#### Blastoids

Blastoids were made using a protocol adapted from the protocol described by Kagawa et al.^13^. On day -1, feeder-free naïve hPSCs^76^ were dissociated using Accutase (5 minutes, 37°C). Dissociation was stopped with DMEM/F12 or N2B27, and cells were spun down (5 minutes, 200 rcf). The pellet was resuspended in N2B27 medium supplemented with 10 µM Y-27632 and 0.3% BSA (Sigma-Aldrich, A7979-50ML). 500 µL of N2B27Y medium containing about 120,000 cells was seeded per well of an anti-adherent solution (Stem Cell Technologies, 07010)-coated AggrewellTM 400 plate (Stem Cell Technologies, 34415). Between day 0 and day 3, blastoids were cultured in PALLY medium consisting of N2B27 medium supplemented with 1 µM PD0325901, 1 µM A83-01 (PeproTech, 9094360), 500 nM lysophosphatidic acid (LPA) (Torcis, 3854), 10 ng/ml human LIF (PeproTech, 300-05-5UG), 10 μM Y-27632, and 0.3% BSA. From day 3 onward, blastoids were cultured for 2 days in LY medium consisting of N2B27 medium supplemented with 500 nM LPA, 10 µM Y-27632 and 0.3% BSA. Blastoids were incubated at 37 °C in hypoxic conditions (5% O2, 5% CO2), and the medium was changed daily. Blastoids in Fig. 6 and Supplementary Fig. 4c, Supplementary Fig. 9, and Supplementary Fig. 10 were generated using 1 µM LPA.

#### SMARCA2 and SMARCA4 degradation

ACBI1 (MedChemExpress, HY-128359) was used to target SMARCA2/4 for degradation. For this, 50 nM ACBI1 was used, which was added fresh to the culture media with every media change. An equal volume of DMSO (Sigma, D2650-100ML) was used as a control. In every experiment, ACBI1 was added to the media at the moment of seeding the cells (day 0 for PXGL, N2B27, nTE, and day -1 for blastoids). For degradation of SMARCA2, ACBI2 (MedChemExpress, HY-151623) was used, similar to ACBI1. Degradation of SWI/SNF ATPases was assessed in most experiments by immunofluorescence.

#### Cell counting and viability quantification

Cell counting and viability assessment were performed using the Countess II (Thermo Fisher). In short, naïve hPSCs were dissociated into single cells using Accutase (Sigma, A6964-500ML) (5 minutes at 37°C). Dissociation was stopped with DMEM/F12 or N2B27, and cells were spun down (5 minutes, 200 rcf), and the pellet was resuspended in N2B27 medium. Cells were diluted 1:1 in 0.4% Trypan Blue. Cell count and viability were subsequently measured using the default settings.

### Embryo culture

#### Consent

Human cryopreserved embryos were used following approval by the local ethical committee of UZ Brussel, Belgium (BUN1432021000539) and the Belgian Federal Committee for research on human embryos (Reference: ADV090). After informed written consent and 5 years of cryopreservation, human surplus embryos were donated to research from patients who completed their fertility treatment.

#### Culture of human embryos

Embryos were warmed and cultured as previously described^77,78^. In short, 3 dpf (8-12 cell stage) or 5 dpf embryos were warmed using the Vit Kit-Thaw kit (90137-SO, Irvine Scientific) following the manufacturer’s instructions. Embryos were given a 2-hour recovery period in equilibrated 25 μl droplets of blastocyst media (83060010, ORIGIO® Sequential Blast™, with phenol red), covered in OVOIL^TM^ (10174, VitroLife) at 37 °C, 5% O2 and 6% CO2. Only embryos that resumed their development after cryopreservation were used in experiments. Embryos were next cultured in a 25 μl droplet of blastocyst media, supplemented with 50 nM ACBI1 or an equal volume of DMSO (1:1000), covered in OVOIL^TM^ at 37 °C, 5% O2 and 6% CO2 in a G185 flat bed incubator (KSystems, CooperSurgical) for 24 **(Fig. 3h)** or 48 **(Fig. 3a)** hours. For the embryos used in **Fig. 1**, 3 dpf and 5 dpf embryos were warmed and cultured until 5 dpf and 6 dpf, respectively, and warmed and cultured in the procedure described above, without ACBI1 or DMSO.

### Immunofluorescence

#### Cultured cells

Immunofluorescence staining was performed as previously described^5,43^. Cells were grown on glass coverslips before collection, washed with 1X PBS (Gibco, 10010015), then fixed in 4% formaldehyde (Life tech, 28908) diluted in 1X PBS, and permeabilized with 1X PBS containing 0.5% Triton X-100 (Sigma, 101597833) (5 minutes) followed by two washes with 1X PBS containing 0.2% Tween-20 (VWR, 0777-1L). Cells were either used immediately or stored at 4°C in 0.2% Tween-20 in 1X PBS. Primary and secondary antibodies (**Supplementary Table 6**) were diluted in a blocking buffer made of 0.2% Tween-20, 5% normal donkey serum (Sigma, S30-100ML), and 2% fish skin gelatin. After incubation with primary antibodies overnight at 4°C, cells were washed three times with 0.2% Tween-20 and incubated with the appropriate fluorophore-labelled secondary antibodies for 60 minutes in the dark at room temperature. Cells were washed with 0.2% Tween-20, 1:50,000 DAPI (1 mg/ml), and washed again with 0.2% Tween-20 in 1X PBS, before mounting in ProLong Gold antifade reagent with DAPI (Invitrogen, P36931). All washes were performed for 5 minutes. Slides were then cured overnight in the dark at room temperature before imaging.

For **Supplementary Fig. 3e**, to improve cell attachment for nTE conversion on coverslips, coverslips were first coated with 0.1% Gelatin (Sigma, G2500, 500G) in 100% Milli-Q water for at least 20 minutes at 37°C. Gelatin-coated coverslips were then coated overnight with 1.5 μg/ml iMatrix Silk Laminin 511 at 37°C. Culture conditions were the same as described above. Samples were collected every day of differentiation.

#### Blastoids

Blastoids were transferred to a well plate (Corning, 3516 or 3513) and were fixed in 1X PBS containing 4% formaldehyde for 30 minutes at room temperature. The blastoids were washed three times with 1X PBS and kept at 4°C until the primary staining was performed. Blastoids were picked and transferred to a Nunc MicroWell 96-Well Microplate (Thermo Scientific, 243656) or round-bottom 96-well plates (Corning, 3370), where they were blocked and permeabilised for 60 minutes at room temperature in blocking and permeabilisation buffer (BPB) consisting of 0.3% Triton-X100 and 10% normal donkey serum in 1X PBS. The primary antibodies were diluted in BPB, and the blastoids were incubated overnight at 4°C with antibodies **(Supplementary Table 6)**. Three washes were performed with 0.1% Triton-X100 in 1X PBS for 5 minutes at room temperature. Secondary antibodies were diluted in 0.1% Triton-X100 in 1X PBS supplemented with 0.01% DAPI (5 mg/ml) and incubated in the dark for 1 hour at room temperature **(Supplementary Table 6)**. Three washes were performed with 0.1% Triton-X100 for 5 minutes at room temperature. Blastoids were transferred to a 15 µ-slide (Ibidi, 81507) and imaged in 1X PBS.

#### Human embryos

Embryos were fixed in 4% PFA in 1X PBS (J61899.AK, Thermo Scientific Chemicals) for 10 minutes at room temperature to cross-link proteins and preserve cellular integrity. After fixation, embryos were washed in 2% BSA (A2153, Sigma-Aldrich) in 1X PBS (10010023, Gibco^TM^) for 5 minutes and were further permeabilised to increase antibody binding efficiency in 0.1% Triton^TM^ X-100 (Sigma-Aldrich) in 1X PBS for 20 minutes at room temperature, with agitation. To decrease non-specific antibody binding, embryos were transferred to 10% FBS-2% BSA in 1X PBS buffer for blocking incubation for 1 hour at room temperature. Embryos were then incubated in the primary antibody, diluted in 10% FBS-2% BSA in 1X PBS overnight at 4 °C. Upon overnight incubation, embryos were washed in 2% BSA-1X-PBS (3 ✕ 10 minutes) and were then incubated in the secondary antibody, diluted in 10% FBS-2% BSA in 1X PBS for 1 hour. From this step forward, embryos were incubated in the dark at room temperature with agitation. For nuclear staining, embryos were further incubated in Hoechst 33342 (H3570, Invitrogen^TM^), diluted in 10% FBS-2% BSA in 1X PBS for 20 minutes. Finally, embryos were washed in 2% BSA in 1X PBS (3 ✕ 5 minutes) and transferred to 10 μl droplets of 2% BSA in 1X PBS on ibidi μ-slide 8-well (80826) or the ibidi μ-slide 15-well 3D (81507) for imaging.

### Microscopy

#### Brightfield

Brightfield images of stem cells and blastoids were taken using a Nikon Eclipse Ti2 microscope or the Thermo Fisher EVOS M3000. Images were analysed using the FIJI ImageJ software^79^.

#### Confocal microscopy

Cultured cells and blastoids were imaged using the Nikon TiE A1R inverted confocal microscope with NIS-Elements AR software. *Z*-stacks were captured through the entire depth of the blastoids or 2D images of cultured cells, at a magnification of 20X, with one to three micrometres distance per picture. Multiple blastoids were captured per image. Multi-channel images were processed using ImageJ and IMARIS. Images were taken at the Bioimaging core of the VIB and KU Leuven. Embryos were visualised for fluorescence using a confocal microscope (LSM800 equipped with Airyscan, ZEISS) at 40X magnification. For whole-embryo visualisation to reconstruct the 3D structure of the embryo, z-stack scans were employed. Images were analysed using FIJI ImageJ and IMARIS. For embryos, Z-stacks were processed with the Zen-Blue software (ZEISS) and image sequences were exported and analysed for cell counts with FIJI ImageJ Suite, including the number of nuclei.

#### Epifluorescence microscopy

Immunofluorescence pictures of cultured cells were measured using the Zeiss LSM 780 confocal microscope, which was converted to function as an epifluorescence microscope by the VIB Bioimaging core in Leuven. Images were analysed using the FIJI ImageJ software.

### Image Analysis

#### ImageJ

Immunofluorescence images were processed using FIJI ImageJ (version 1.54g), and each channel was manually adjusted for brightness and contrast. The same settings per channel were used for each staining within each experiment. The Despeckle function was used to remove noise from some of the images that are used in the panels.

The measurement function of FIJI ImageJ was used to measure the diameter of the blastoids. For this, brightfield images were taken daily. The horizontal axis was used to measure the diameter of each blastoid.

Fluorescence intensities of SMARCA2, SMARCA4 and SOX2 in human embryos were measured using the 3D Fiji/ImageJ suite in FIJI software as described previously^72,80^. Briefly, the nuclei of immunolabelled embryos were segmented automatically based on the signal of Hoechst-33342 and the marker of interest. Fluorescent intensities were measured within the nuclei.

For **Supplementary Fig. 3e-g**, immunofluorescence images of nTE differentiation on coverslips were analysed with FIJI ImageJ. Colonies were identified using the DAPI channel by applying an intensity threshold, after which the Analyse Particles function was used to extract mean fluorescence intensity values of cells for each marker channel. DAPI intensity was used to normalise the signal across cells. Thresholds to classify GATA2-positive and NANOG-positive cells were determined based on the signals in WT nTE differentiation and in naïve hPSCs, respectively. The same settings were used per channel for each staining within each experiment.

#### IMARIS

For images in **Fig. 2** and **Supplementary Fig. 1d** and **Supplementary Fig. 4b**, the analysis of SMARCA2 and SMARCA4 intensity, IMARIS (Version 10) was used to analyse immunofluorescence images. Cells were detected using the spot-detection tool on the DAPI channel. This function was used for quantification of blastoid cell number and used as a basis for cell lineage and signal intensity quantification. For each image containing multiple blastoids, the machine learning tool was used to assign the detected cells to a specific cell lineage (EPI/TE/HYPO), which was manually curated based on location and marker expression. For signal intensity analysis, DAPI intensity was used to normalise the loss of signal throughout the z-stack.

For images in **Fig. 3**, **Supplementary Fig. 4i-j**, **Supplementary Fig. 6**, and **Supplementary Fig. 9**, IMARIS was used to analyse immunofluorescence images. Cells were detected using the spot-detection tool per channel to detect marker-positive cells. For each image containing multiple blastoids, a signal intensity value was used in the control condition, which was manually curated based on location and marker expression of the cell. For signal intensity analysis, DAPI intensity was used to normalise the loss of signal throughout the z-stack.

#### Zen Blue

For embryos, Z-stacks were processed with the Zen Blue software (ZEISS) and image sequences were exported and analysed for cell counts with FIJI ImageJ Suite, including the number of nuclei. The diameters of embryos (with and without zona pellucida) were measured using the ‘measure’ function in the image analysis module. Embryonic lineage numbers (TE, EPI, and HYPO) were assessed manually. A nucleus was considered positive if the signal was visibly higher than the background.

### Flow cytometry

#### Cells were dissociated with Accutase or 0.25% Trypsin-EDTA (nTE)

Fluorophore-conjugated antibodies were diluted in 0.1% BSA in 1X PBS (FACS buffer) for 20 minutes at room temperature, then washed with FACS buffer, and fixed in 4% formaldehyde. Cells were passed through a 40 µm cell strainer (Corning 352340). For viability staining, 1:300 Live/Death Zombie Aqua (Biolegend, 423102) in 1X PBS was used before antibody staining. 10,000 cells were acquired using the BD FACSymphony A5 or the BD FACSCanto II AIG and analysed using the FlowJo software (version 10.10, BD Bioscience). Single-stained and unstained controls were used for compensation and gating.

#### RNA extraction using TRIzol

Cells were collected on days 1, 2, and 3 of nTE conversion using TrypleE Express (Gibco, 12605-010) or 0.25% Trypsin-EDTA (1X) (Gibco, 25200-056) for 5-10 minutes to achieve single-cell dissociation. Dissociation was stopped by diluting the samples in N2B27. Cells were centrifuged (5 minutes, 400 rcf), and cell pellets were stored at -80°C until further use. The cell pellet was lysed using TRIzol (Invitrogen, 15596024) for 5 minutes. Chloroform (VWR Chemicals, 67-66-3) was added to the samples, mixed, incubated for 3 minutes, and centrifuged at 10,000 g for 5 minutes at 4°C. The aqueous phase was transferred to a new tube. Isopropanol (VWR Chemicals, 67-63-0) was added, and the samples were incubated for 10 minutes before centrifugation at 10,000 g for 10 minutes at 4°C. The supernatants were discarded, and the cell pellet was resuspended in 75% ethanol (VWR Chemicals, 64-17-5) in ultrapure water (Thermo, 10977035). Samples were centrifuged at 10,000 g for 5 minutes at 4°C. The supernatants were removed, and the pellet was air-dried for 10 minutes. The cell pellet was resuspended in 30 μl ultrapure water. RNA concentration was determined using the NanoDrop (Thermo Scientific).

#### cDNA synthesis

The RNA was diluted in ultrapure water to obtain 2 μg in 10.6 μl. To the diluted DNA, a MasterMix was added consisting of 2.5 ng/μl Random hexamers (Thermo Fisher, S0142), 5 μg/ml Oligo(dt) 12-18 (Invitrogen, 18418012), 500 μM dNTP mix (Invitrogen, 10297-018), 1:10 M-MLV reverse transcriptase (homemade), RNaseOUT (Invitrogen, 10777019), 1X First Strand Buffer (prepared in-house), and 10 mM DTT (Invitrogen, 707265ML). cDNA synthesis was performed using the thermal cycler (Thermo Fisher), with a program of 10 minutes at 25°C, 50 minutes at 42°C, 15 minutes at 70°C, and 4°C. Samples were stored at -20°C until further use.

#### qPCR

Quantitative PCR was performed using a 384-well plate in the ABI ViiA7 real-time PCR system (Applied Biosystems). cDNA was diluted in ultrapure water, and 1.25 ng cDNA was used for qPCR, together with 45% Platinum SYBR Green platinum mix (Thermo Fisher, 11733046), 0.02% ROX reference dye (Thermo Fisher, 12223012), and 0.25 μM of the forward and reverse primers **(Supplementary Table 6)**. The PCR plate was centrifuged for 1 minute at 1000 rpm and subsequently loaded into the ViiA7.

#### RT-qPCR analysis

The ΔCt was calculated using GAPDH as the reference gene^58^. The ΔΔCt was calculated for pluripotency markers using the ΔCt of naïve hPSC as the control sample, and for TE markers using the ΔCt of D3 nTE WT as the control sample. The fold change was then calculated as follows: 2^(-ΔΔCt).

### Single-nucleus multiomic RNA plus ATAC sequencing

#### Sample preparation and nuclei isolation

Blastoids were generated as described above from SIGi001-A naïve hPSCs. Half of the blastoids were fixed for validation, and the other half were frozen in freezing media consisting of 5% DMSO in N2B27. There was no selection of blastoids for library preparation. Blastoids were thawed and diluted in warm N2B27. After centrifugation for 5 minutes at 500 rcf, the pellet was resuspended in a lysis buffer containing 10 mM Tris-HCl (pH 7.4, Sigma, T2194-100ML), 10 mM NaCl (Invitrogen, AM9760G), 3 mM MgCl_2_ (Invitrogen, AM9530G), 0.1 % Igepal, 1% BSA, 1U/µl RNase Inhibitor (10X Genomics, 2000565) in nuclease-free water (Thermo scientific, AM9937). The samples were centrifuged through 20 and 10 µm filters (Imtec, 43-10010-60/43-10020-60) for 5 minutes at 500 rcf. The pellet was then resuspended in 0.1% lysis buffer containing 10 mM Tris-HCl (pH 7.4, Sigma, T2194-100ML), 10 mM NaCl, 3 mM MgCl_2_, 0.01 % Igepal, 1% BSA, 1U/µl RNase Inhibitor, 0.01% Tween-20 (Bio-rad, 1662404), 0.002% Digitonin (Merck, 300410-1GM), and 1 mM Dithiothreitol (DTT, Sigma, 646563-10X.5ml) in nuclease free water. After an incubation of 1 minute on ice, protector buffer consisting of 10 mM Tris-HCl (pH 7.4, Sigma, T2194-100ML), 10 mM NaCl, 3 mM MgCl_2_, 1% BSA, 1U/µl RNase Inhibitor, 0.1% Tween-20, and 1 mM DTT in nuclease-free water was added. The samples were then centrifuged through a 10 µm filter for 5 minutes at 500 rcf, and the pellet was resuspended in diluted nuclei buffer, consisting of 1X Nuclei buffer (10X Genomics, 2000207), 1 mM DTT, and 1 U/µl RNase Inhibitor in nuclease-free water. Nuclei were counted using the Luna counter by adding 9 µl of the nuclei suspension with 1 µl of AO/PI dye. The nuclei were diluted to 1900 nuclei/µl in diluted nuclei buffer and used for library preparation.

#### Library preparation

Library preparation was performed according to the 10X Genomics Chromium Next GEM Single Cell Multiome ATAC + Gene Expression user guide (10X Genomics, CG000338 Rev F). 3000 nuclei were targeted. In short, nuclei were isolated as described above and transposed. Gel beads-in-emulsion (GEMs) were generated to generate barcoded transposed DNA and cDNA. The pre-amplified samples were then split into two to generate the ATAC and gene expression libraries.

For the ATAC library, 8 cycles were used for the Sample Index PCR. For the gene expression library, 7 cycles were used during cDNA amplification, and 14 cycles were used for the Sample Index PCR. DNA concentrations were determined using the Qubit (Thermofisher, Q32854). Quality control and library concentrations were determined using the BioAnalyzer (Agilent) or the Tapestation (Agilent, Tapestation 4150) using the High Sensitivity D5000 ScreenTape assay (Agilent, 5067-5592/3). Molarity calculations were performed using the formula: 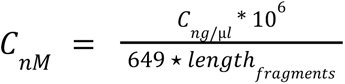. After library generation, the libraries were pooled per library type, molarity was measured, and the samples were diluted to 2 nM for sequencing.

#### Sequencing

Libraries were sequenced at 750 pM using the NextSeq2000, using the P1-100 cycles flowcell with XLEAP chemistry (Illumina). The cycles for Read1, i7, i5, and Read2 were 50,8,25,49 and 28,10,10,90 for the ATAC and gene expression libraries, respectively. We received sequencing data of 6022 cells (DMSO: 2405, ACBI1: 3617). For the DMSO gene expression library, we obtained 214,490,747 read pairs with 89,185.34 mean raw reads per cell, over 1746 median UMIs per cell, detecting a median of 945 genes per cell. For the ACBI1 gene expression library, we obtained 258,266,492 read pairs with 71,403.51 mean raw reads per cell, over 1497 median UMIs per cell, detecting a median of 661 genes per cell. For the DMSO ATAC library, we obtained 259,923,092 read pairs with 108,076.13 mean raw read pairs per cell for a median of 5728 high-quality fragments per cell. For the ACBI1 ATAC library, we obtained 220,116,891 read pairs with 60,856.20 mean raw read pairs per cell for a median of 6587 high-quality fragments per cell.

After filtering (described below), we obtained gene expression data from 4556 cells (DMSO: 1571, ACBI1: 2985) and Assay for Transposase-Accessible Chromatin sequencing (ATAC) data from 3431 cells (DMSO: 1440, ACBI1: 1991), of which 2623 cells had both gene expression and ATAC data (DMSO: 959, ACBI1: 1664).

### PIPseq single-nucleus transcriptome sequencing

#### Sample preparation and nuclei isolation

Blastoids were generated as described above from SIGi001-A naïve hPSCs and frozen in freezing media consisting of 5% DMSO in N2B27. There was no selection of blastoids for library preparation. Blastoids were thawed and diluted in warm N2B27. After centrifugation for 5 minutes at 500 rcf, the pellet was resuspended in a lysis buffer containing 10 mM Tris-HCl (pH 7.4, Sigma, T2194-100ML), 10 mM NaCl (Invitrogen, AM9760G), 3 mM MgCl2 (Invitrogen, AM9530G), 0.1 % Igepal, 1% BSA, 0.8 U/µl Protector RNase Inhibitor (Sigma, 03335399001) in nuclease-free water (Thermo scientific, AM9937). The samples were centrifuged through a 20 and 10 µm filter (Imtec, 43-10010-60/43-10020-60), for 5 minutes at 500 rcf. The pellet was then resuspended in 0.1% lysis buffer containing 10 mM Tris-HCl (pH 7.4, Sigma, T2194-100ML), 10 mM NaCl, 3 mM MgCl2, 0.01 % Igepal, 1% BSA, 0.8 U/µl

Protector RNase Inhibitor, 0.01% Tween-20 (Bio-rad, 1662404), 0.002% Digitonin (Merck, 300410-1GM), and 1 mM Dithiothreitol (DTT, Sigma, 646563-10X.5ml) in nuclease free water. After incubation for 1 minute on ice, protector buffer consisting of 10 mM Tris-HCl (pH 7.4, Sigma, T2194-100ML), 10 mM NaCl, 3 mM MgCl2, 1% BSA, 0.8 U/µl RNase Inhibitor, 0.1% Tween-20, and 1 mM DTT in nuclease-free water was added. The samples were then centrifuged for 5 minutes at 500 rcf, and the pellet was resuspended in nuclei suspension buffer (NSB), consisting of nuclei suspension buffer (NSB, Illumina Single Cell 3’ RNA Capture, T2, Box 3), 1% BSA (25% BSA, Illumina Single Cell 3’ RNA Capture, T2, Box 3), and 0.8 U/µl SUPERase-In RNase Inhibitor (Invitrogen, AM2694) in nuclease-free water to wash the nuclei. Nuclei were centrifuged for 5 min at 500 g through a 10 µm filter, and the pellet was resuspended in NSB. Nuclei were counted using the Luna counter by adding 9 µl of the nuclei suspension with 1 µl of AO/PI dye. The nuclei were diluted to a target of 1250 nuclei/µl in NSB and used for library preparation.

#### Library preparation

Library preparation was performed according to the Illumina Single Cell 3’ RNA prep T2 protocol (Illumina, 200063178 v01). 2000 nuclei were targeted. In short, nuclei were isolated as described above. Particle-templated instant partitions (PIPs) were generated using the PIPseq Vortex Mixer (Fluent BioSciences, FBS-SCR-DVM), and the nuclei were lysed using the Prog. B settings of the PIPseq Dry Bath (Fluent BioSciences, FBS-SCR-PBD).

PIPs were isolated, volume-normalised, and the mRNA was reverse-transcribed to cDNA. PIPs were washed, and the cDNA was amplified. After cDNA amplification, the cDNA was isolated, and quality control was performed after an additional amplification with 13 cycles using the Tapestation (Agilent) High Sensitivity D5000 ScreenTape assay (Agilent, 5067-5592/3). cDNA was fragmented, end-repaired, and A-tailed before adapter ligation. DNA concentrations were determined using the Qubit (Thermofisher, Q32854). After clean-up of the ligated fragment, sample indexes (Illumina, UDI Mix Strip, 20137528) were added, and the library was amplified using 16 cycles. Quality control was performed using the Tapestation (Agilent, Tapestation 4150) with the High Sensitivity D5000 ScreenTape assay (Agilent, 5067-5592/3). Libraries were pooled using equal volumes with equal molarities, and the library was diluted to 2 nM for sequencing.

#### Sequencing

Libraries were sequenced at 550 pM using the NextSeq2000, using the P3-100 cycles flowcell with XLEAP chemistry (Illumina). The cycles for Read1, i7, i5, and Read2 were 45, 10, 10, and 72. BCL files were converted to FASTQ files using BCL Convert (Illumina, version 4.5.4). FASTQ files were subsequently converted to counts with PIPseeker (Fluent BioSciences, version 3.3.0) using the *pipseeker-gex-reference-GRCh38-2022.04* reference genome. PIPseeker’s Sensitivity 5 outputs were used for further analysis **(Supplementary Table 7)**.

### Bioinformatics

All code will be made available on GitHub: https://www.github.com/pasquelab. ChatGPT (version 4, 5, unpaid), Claude (Sonnet 4.6, 5, unpaid), and M365 Copilot (GPT-4) have been used for troubleshooting during data analysis.

#### Data preparation

The pipeline of Cellranger ARC version 2.0.0 was used to convert raw BCL files into FASTQ files, and subsequently into counts. For the reference genome, Human reference cellranger arc GRCh38 2020 A version 2.0.0 was used.

#### RNA sequencing analysis of published datasets

Bulk RNA-seq data analysis of the nTE differentiation was performed based on the R script provided by GEO2R, using DESeq2 version 1.44.0 (GSE144994)^6^. For this, we selected the data of naïve hPSCs and days one to three of nTE differentiation. Genes were filtered out with fewer than 10 counts. Counts were log10-normalised.

Single-cell RNA-seq data analysis of human blastoids was performed as described previously^5^. Data were log-normalised and scaled using Seurat version 4.2.0^81^. For this analysis, we used the original FACS-sorted annotation per timepoint from the original publication^13^.

For SMARCA2/4 RNA expression analysis, the human embryonic reference tool version 1.1.1.17 was used^17^.

#### Mass spectrometry of stem cell models

Stem cell mass spectrometry samples (naïve hPSCs, primed hPSCs, hTSCs) presented in Girard et al.^52^ were re-analysed with a DIA acquisition using the nanoliquid chromatography-tandem mass spectrometry analysis settings of Onfray et al.^82^. The different DIA raw files were analysed with Spectronaut software version 19 (Biognosys, Schlieren, Switzerland) using direct DIA default settings. Data were searched against the UniProt KB Human database (20,593 sequences, downloaded in June 2023), with trypsin/P as the protease with up to one missed cleavage. To account for post-translational modifications and chemical labelling settings, carbamidomethylation of cysteine residues was defined as a fixed modification, and methionine oxidation and acetylation of Lysines and acetylation of protein N-termini were defined as variable modifications. The files were analysed with Spectronaut using default settings except for quantification, and allowed quantification of the precursors, peptides and proteins. Quantifications were performed using the mean precursor quantity and then the sum peptide quantity. The results were filtered by a 1% FDR at the precursor, peptide and protein level using a target-decoy approach, which corresponds to a Q value % 0.01^83^. Quantification data were then normalised by Spectronaut software to take into account the overall acquisition heterogeneity between samples. Given the number of samples analysed (less than 500 individuals), the type of data normalisation carried out for the whole dataset was a local regression normalisation described by Callister et al. 2006 ^84^. Protein expression levels were then log2-transformed. Dotplots were processed using the package ggplot2.

#### Mass spectrometry of human embryos

Human embryo mass spectrometry samples (oocyte, 1-cell, 4-cells, 8-cells, morula, and blastocyst) presented in Zhu et al.^53^ were used to plot the expression of SMARCA2 and SMARCA4. In brief, protein quantification was performed on Spectronaut V18 with default settings except for the normalisation. Protein expression levels were first normalised within each sample using the upper quantile expression value, then log2-transformed and normalised on quality control samples. Dotplots were processed using the package ggplot2.

#### Multiome Analysis

SCANPY version 1.11.2^85^ was used to analyse gene expression data. Cells were filtered by; n_genes_by_counts < 12,000, total_counts < 75,000, percentage mitochondrial genes (pct_counts_mt) < 5, min_genes = 250, min_cells = 3. Counts were normalised to a target sum of 1e4 and log-transformed. Counts were regressed to total_counts and pct_counts_mt, and scaled. Dimensionality reduction was performed using 20 neighbours and 12 principal components. UMAP clustering was performed using the Leiden algorithm with a resolution of 0.1. The clusters were manually annotated. Further analysis was performed on the data stratified by both treatment and cluster.

PyCisTopic version 2.0a0^22,59^ was used to analyse the ATAC data. The single-nucleus ATAC data were pseudobulked based on the treatment-cluster annotation. Peaks were called from the pseudobulk profiles using pyCisTopics’ peak_calling, and consensus peaks were created. Cells were filtered based on the automatic thresholds. Doublets were removed using Scrublet. We performed an analysis on 20 topics. UMAP clustering was performed using 10 neighbours and a resolution of 0.1, and clusters were annotated based on the overlap with the gene expression clustering. Topics were binarised, differentially accessible regions were assessed, and gene activity was calculated. Coverage plots were made using Seurat/Signac’s BigwigTrack with the bigwigs from the pseudobulked profiles.

Genomic region annotation of ATAC topics, cummulated peaks summits per sample, and differentially accessible regions were performed using ChIPseeker version 1.42.0. For topic analysis, the top 3000 regions per topic were used. In short, BED files were converted to a GRanges object using the GRanges command. The peaks were subsequently annotated using annotatePeak, using *TxDb.Hsapiens.UCSC.hg38.knownGene* for TxDb and annoDb as org.Hs.eg.db. The promoter was defined as the region within 3000 bp of the TSS.

For integrated multiome analysis, the SCENIC+ version 1.0a1 was used^22^. For this, a custom pyCisTarget database was generated using pyCisTarget version 1.0a2^22^, with the human reference genome hg38 from UCSC. After running SCENIC+ for 10 iterations, the direct and extended gene-based AUCs were concatenated per iteration, and UMAP dimensional reduction was performed. Per iteration, the eRegulon specificity score (eRSS) was calculated based on the direct eRegulons. We then averaged the eRSS per eRegulon name over the 10 iterations and removed the regulons occurring less than 5 times.

For the integration of our gene expression data with the human embryo data, the Early Embryogenesis Projection Tool (v2.1.2) was used^17^. To obtain the count matrix, Seurat version 5.3.0^86^ was used, and cells were filtered based on the barcodes that passed filtering in SCANPY. A correlation filtering of 0.5 was used for the integration.

Some downstream analyses, including dot plots, module score calculations, and motif accessibility, were performed using Seurat version 5.1.0, Signac version 1.14.0 and ChromVar version 1.28.0. For this, a Seurat object was created, and cells were filtered based on the barcodes that passed the SCENIC+ pipeline, described above. The gene expression dataset was normalised by SCTransform, and the ATAC dataset was normalised using RunTFIDF. This Seurat object was then used to make the DotPlots for gene expression. Module scores were calculated using the AddModuleScore function with genes provided by Meisterman et al. (gene modules)^49^, the MSigDB (M5902, M5901, M11558, M14297, M13413, M14052, M17089, and M5936), and the identified target genes of SCENIC+.

Motif accessibility analysis was performed using Seurat version 5.1.0, Signac version 1.14.0 and ChromVar version 1.28.0 with JASPAR2020 as the TF motif database. Accessibility was calculated using the getMatrixSet, CreateMotifMatrix, and CreateMotifObject, followed by the RunChomVar command, on the same cells that previously passed the SCENIC+ analysis with the same annotation. For this, the BSgenome.Hsapiens.UCSC.hg38 genome was used.

Correlation of TF expression and TF motif accessibility was performed by filtering the Seurat-normalised counts and the ChromVar accessibility score above for the same genes. Panda’s corrwith was used to calculate the correlation score between the expression of a TF and its corresponding motif accessibility. Slopes were calculated in R, using the *lm* function per Sample.

Profile plots were generated using DeepTools version 3.5.6^87^. For this, genome region scores were calculated using *computeMatrix reference-point*, pyCisTopic’s pseudobulked bigwig files were used as input. The region was set at the centre of the BED files (from published datasets^42,60,64^, DARs, SCENIC+ predicted enhancer regions, and the Eukaryotic Promoter Database^88^) plus/minus 3000 bp.

Differential gene expression analysis was performed using Seurat’s *FindMarkers*. For downstream analysis, genes were filtered using p_val_adj < 0.05 and avg_log2FC > 0.5, or < -0.5.

#### PIPseq analysis

Seurat version 5.5.0^86^ was used to analyse the PIPseq single-nucleus transcriptome data. Cells were filtered using; nFeature_RNA > 200, nFeature_RNA < 11000, percent.mt < 5, and nCount_RNA < 50000. Counts were normalised using Seurat’s *SCTransform*. Dimensionality reduction was performed using 20 neighbours and 15 principal components. UMAP clustering was performed using the Leiden algorithm with a resolution of 0.2. The clusters were manually annotated based on marker gene expression. Further analysis was performed on the data stratified by both sample and cell type annotation.

Trajectory analysis was performed based on CellRank2 (version 2.3.2)^65^. For this, the transcriptome data was reanalysed using SCANPY (version 1.12.3) (Wolf et al. 2018). Cells were filtered to include the cells that passed the filtering above. Samples were combined into three objects based on the day 3 sample (Trajectory 1: day 3 DMSO + day 5 DMSO + day 5 ACBI1 3:5. Trajectory 2: day 3 ACBI1 -1:0 + day 5 ACBI1 -1:0. Trajectory 3: day 3 ACBI1 + day 5 ACBI1 + day 5 ACBI1 -1:3). Counts were normalised to a target sum of 1e4 and log-transformed. Dimensionality reduction was performed using 30 neighbours and 11 principal components. Pseudotime analysis was performed per trajectory with Palantir (version 1.4.5)^89^, using SCANPY. For this, a cell from the day 3-EPI cluster was chosen as the initial state, and 500 waypoints were used. The SCANPY objects were concatenated with Velocyto objects^90^, using scVelo (version 0.3.4)^91^. For this, sorted BAM files were generated using PIPseeker, and Velocyto objects were generated using Velocyto.py (version 0.17.17), and samples were re-normalised and log-transformed. Subsequently, CellRank was performed on the scVelo object, and a transition matrix was computed on the pseudotime kernel. Downstream analysis was performed using the GPCCA estimators. Macrostates were computed using 8, 5, and 6 states from trajectories 1-3, respectively. From these macrostates, initial and terminal states were manually set and used to compute fate probabilities.

Differential gene expression analysis was performed using Seurat’s *FindAllMarkers*, and pairwise comparisons were performed using Seurat’s *FindMarkers*. For downstream analysis, genes were filtered using p_val_adj < 0.05 and avg_log2FC > 0.5 or < -0.5.

The PIPseq and multiome transcriptomes were integrated using Seurat’s pipeline for SCTransform-based normalised integration. In short, PIPseq and multiome counts were normalised using SCTransform. The top 3000 features were selected for integration using *SelectIntegrationFeatures*. Seurat objects were prepared for integration using *PrepSCTIntegration*. Integration anchors were found using the *FindIntegrationAnchors* function, using SCT as the normalisation method. Finally, the PIPseq and multiomic datasets were integrated using *IntegrateData*, with SCT as the normalisation method. After integration, dimensionality reduction was performed on the first 30 principal components.

### Statistics

The Statistical Consulting group of Biomedical Sciences Leuven was consulted for parts of the statistical approach in this study. All statistical output can be found in **Source Data**. Statistics were performed in R (version 4.4.1).

For **Fig. 2c**, per experiment, the change in DAPI-corrected intensity was calculated per condition. For this, first the median DAPI-corrected intensity was calculated. Then, the median Intensity of the DMSO condition was subtracted, and this difference was divided by the median Intensity of the DMSO condition. To test whether this value was decreased, we performed a one-sided one-sample t-test using the *t.test* function in R, and set the alternative hypothesis to whether the value was less than 0. The p-values were adjusted using the Bonferroni method. N = 5 experiments.

For **Fig. 2e**, a step-wise approach was used. First, the mean diameter per data point was log-transformed, giving us the log of the mean size (LMS) per day and sample. We then performed the lm function in R on the LMS as a function of time, treatment, and the interaction between time and treatment. Here we showed that time and the interaction between time and treatment, but not treatment alone, were significant. Subsequently, a two-sided Wilcoxon rank-sum exact test was performed using the *wilcox.test* function on the mean size, stratified by day, to determine whether on each day the size was different. The p-values are Bonferroni-corrected. N = 5 experiments.

For **Fig. 2h**, equal variance was tested using the var.test function, and normality was tested using the *shapiro.test* function. As the conditions had equal variance and the data were normally distributed, a two-sided two-sample t-test was performed using the *t.test* function to determine significance. N = 7 experiments.

For **Fig. 2i**, a step-wise approach was used. First, to test the difference in cell lineage composition, Aitchison’s test^92^ was performed using the *ait.test* function of the Compositional package version 7.3, on the relative composition per lineage. As this test does not allow zeroes, we set all zeroes to 0.0000000001. Next, Pearson’s Chi-squared test was performed using the *chisq.test* function to assess the significance of the cumulative cell number count per lineage and condition. These tests showed that both the composition and the overall cumulative counts were significantly different. Next, Fisher’s Exact Test was performed on the cumulative counts categorised per lineage using the *fisher.test* function to determine the significance per lineage. N = 8 experiments, n = 20423 (DMSO) and 26008 (ACBI1) cells.

For **Fig. 3c, a** two-sided Wilcoxon rank-sum test with continuity correction was performed using the *wilcox.test* function. P-values were corrected using Bonferroni correction. N = 7 embryos per condition.

For **Fig. 3d** and **e**, equal variance was tested using the var.test function, and normality was tested using the *shapiro.test* function. As the conditions had equal variance and the data were normally distributed, a two-sided two-sample t-test was performed using the t.test function. N = 21 (DMSO) and 19 (ACBI1) embryos.

For **Fig. 3g and Supplementary Fig. 5c**, **e**, a two-sided Wilcoxon rank-sum test with continuity correction was performed using the *wilcox.test* function. P-value correction was performed using the Bonferroni method. N = 21 (DMSO) and 19 (ACBI1) embryos.

For **Fig. 3j and Supplementary Fig. 6e**, equal variance was tested using the var.test function, and normality was tested using the *shapiro.test* function. As the conditions had equal variance and the data were normally distributed, a two-sided two-sample t-test was performed using the t.test function. N = 7 (DMSO) and 6 (ACBI1) embryos.

For **Fig. 3k**, per lineage, the Wilcoxon rank-sum test with continuity correction was performed on the lineage ratio per embryo using the *wilcox.test* function. N = 7 (DMSO) and 6 (ACBI1) embryos.

For **Fig. 5h and Supplementary Fig. 8i**, a two-sided paired Wilcoxon signed-rank test with continuity correction was performed using the *wilcox.test* function. N = 300 genes.

For **Supplementary Fig. 1d,** a two-sided Wilcoxon rank sum test with continuity correction was performed using the *wilcox.test* function in R. The p-values were Bonferroni-corrected. N = 3 embryos.

For **Supplementary Fig. 3c**, the Kruskal-Wallis rank sum test was performed using the *kruskal.test* function. The *pairwise.wilcox.test* function was used to perform a two-sided pairwise Wilcoxon rank sum analysis. N = 4 experiments.

For **Supplementary Fig. 3d**, a two-sided pairwise Wilcoxon rank sum exact test was performed using the *pairwise.wilcox.test* function on the fold-change values. P-values were adjusted using the Benjamini & Hochberg correction. N = 5 experiments.

For **Supplementary Fig. 4f**, per experiment, the change in DAPI-corrected intensity was calculated per condition. For this, first the median DAPI-corrected intensity was calculated. Then, the median Intensity of the DMSO condition was subtracted, and this difference was divided by the median Intensity of the DMSO condition. To test whether this value was decreased, we performed a one-sided one-sample t-test using the *t.test* function and set the alternative hypothesis to whether the value was less than 0. Normality was assumed. The p-values were adjusted using the Bonferroni method. N = 3 experiments.

For **Supplementary Fig. 4h**, per day, a one-way ANOVA was performed on the mean size per experiment using the *aov* function, followed by a two-sided pairwise t-test. Normality was assumed. P-values were Bonferroni-corrected. N = 3 experiments.

For **Supplementary Fig. 5f**, a two-sided Wilcoxon rank-sum test with continuity correction was performed using the *wilcox.test* function. P-value correction was performed using the Bonferroni method. N = 14 (DMSO) and 12 (ACBI1) embryos.

For **Supplementary Fig. 5h**, a two-sided Wilcoxon rank-sum test with continuity correction was performed using the wilcox.test function. N = 7 embryos per condition.

For **Supplementary Fig. 6f, g**, per lineage, equal variance and normality were tested using the F test (*var.test)* and the Shapiro-Wilk normality test (*shapiro.test)*. This was followed by a two-sided two-sample t-test (Fig. 6g) or a Wilcoxon rank-sum exact test with continuity correction (Fig. 6f) on the mean marker Intensity per lineage per embryo, using the *t.test* or *wilcox.test* Function, respectively. N = 7 (DMSO) and 6 (ACBI1) embryos.

For **Supplementary Fig. 9b**, per experiment, the change in DAPI-corrected intensity was calculated per condition. For this, first the median DAPI-corrected intensity was calculated. Then, the median Intensity of the DMSO condition was subtracted, and this difference was divided by the median Intensity of the DMSO condition. To test whether this value was decreased, we performed a one-sided one-sample t-test using the *t.test* function and set the alternative hypothesis to whether the value was less than 0. Normality was assumed. The p-values were adjusted using the Bonferroni method. N = 3 experiments.

For **Supplementary Fig. 9d**, per day, a one-way ANOVA was performed on the mean size per experiment using the *aov* function, followed by a two-sided pairwise t-test. Normality was assumed. P-values were corrected using the Benjamini & Hochberg method. N = 3 experiments.

## Artificial Intelligence

Perplexity, ChatGPT(versions 4 and 5, unpaid), Claude (Sonnet 4.6, 5, unpaid), and M365 Copilot (GPT-4) were used to improve the language and readability of parts of the manuscript and for troubleshooting during data analysis. The authors take full responsibility for the content of this article.

## Data and code availability

Upon publication of the manuscript, all code will be made available on GitHub: https://www.github.com/pasquelab.

Sequencing data has been uploaded to ENA and GEO and will be made publicly available upon publication of the manuscript.

## Ethical approval statement

Blastoids recapitulate the entire conceptus. However, these are not considered human embryos according to ISSCR guidelines. Our work complies with the current ISSCR 2006 and 2021 guidelines, as well as the 2025 ISSCR guideline update. Work with human embryonic and induced pluripotent stem cells to model early human development, including stem cell-based human embryo models and blastoids, was approved by the UZ/KU Leuven ethics committee (S66595, S64962, S66185, and S68981). Work with mouse embryonic fibroblasts was approved by the UZ/KU Leuven ethics committee (P170/2019). The human embryo experiments have been approved by the local Commission of Medical Ethics of the UZ Brussel (B.U.N. 1432021000539) and the Federal Committee for Medical and Scientific Research on Human Embryos in vitro (AdV090) and by the UZ/KU Leuven ethics committee (S66184). The ethical approval included the warming, culture, small molecule perturbation, fixation, and immunohistochemistry by confocal microscopy, with the aim of studying chromatin modifiers during the first 14 days of human development. We used cryopreserved human embryos donated to research from IVF-ICSI patients (UZ Brussel) after informed consent and the expiration of the 5-year cryopreservation duration.

Gametes have not been used in this study. All experiments involving human embryos were conducted using embryos between 3 and 5 dpf. We used cryopreserved supernumerary human embryos donated to research from IVF-ICSI patients (UZ Brussel) after informed consent. No embryos were created for this study. As the national legislation and institutional guidelines suggest, human embryos were only used after 5 years of cryopreservation and no use, from only patients who had consented prior to their treatment. If required, details on consent forms can be further provided. The experimental endpoint was consistently set at 5 dpf, meaning that embryos were fixed approximately nine days before 14 dpf and the onset of primitive streak formation. The endpoint for the human blastoids was at day 5 of human blastoid formation, equivalent to the 7 dpf stage of human embryo development.

In this research, embryos were pharmacologically treated with small molecule inhibitors. Following fixation, embryos were analysed by immunofluorescence and confocal microscopy using antibodies against key targets, including selected chromatin remodellers and transcription factors. This approach allowed us to assess how perturbation of SMARCA2/4 influenced cell fate specification in the human embryo. Single-cell genomic analyses were not performed on human embryos.

## Contribution

Conceptualisation: S.S.F.A.v.K (Embryo characterisation, Stem cells, Blastoid, Multiomics, Transcriptomics), M.T (Embryo perturbation), H.V.D.V, V.P.;

Data Curation: S.S.F.A.v.K (Embryo characterisation, Stem cells, Blastoid, Multiomics, Transcriptomics), M.T (Embryo perturbation).;

Formal Analysis: S.S.F.A.v.K (Embryo characterisation, Stem cells, Blastoid, Multiomics, Transcriptomics), M.T (Embryo perturbation), J.D.B (Stem cells, Blastoids), E.B (Stem cells, Blastoids), C.L (Multiomics).; O.G. (Proteomics).

Funding: S.S.F.A.v.K (FWO fellowship), H.V.D.V (Embryo), V.P. (Embryo characterisation, Stem cells, Blastoid, Multiomics, Transcriptomics).;

Investigation: S.S.F.A.v.K (Embryo characterisation, Stem cells, Blastoid, Multiomics, Transcriptomics), M.T (Embryo perturbation), J.D.B (Stem cells, blastoids), E.B (Stem cells, Blastoids), M.S (Stem cells), S.K (Stem cells).;

Methodology: S.S.F.A.v.K (Embryo characterisation, Stem cells, Blastoid, Multiomics, Nuclei isolation, Transcriptomics), M.T (Embryo perturbation), M.W (Multiomics), S.P (Multiomics),

J.D.C (Nuclei isolation), A.M (Nuclei isolation), I.S (Nuclei isolation), E.W (Nuclei isolation),

Y.M (Nuclei isolation), H.V.D.V (Embryo), V.P (Stem cells, Blastoid, Multiomics).;

Project Administration: S.S.F.A.v.K, M.T (Embryo perturbation), H.V.D.V (Embryo), V.P (Stem cells, Blastoid, Multiomics, Transcriptomics).;

Resources: S.P (Multiomics), H.V.D.V (Embryo), V.P (Stem cells, Blastoid, Multiomics, Transcriptomics).;

Visualisation: S.S.F.A.v.K (Embryo characterisation, Stem cells, Blastoid, Multiomics, Transcriptomics), M.T (Embryo perturbation), J.D.B (Stem cells, Blastoids), E.B.(Stem cells, Blastoids), O.G.(Proteomics);

Writing, reviewing, Editing of manuscript: S.S.F.A.v.K, M.T, J.D.B, E.B., S.K, L.D, H.V.D.V, V.P

## Acknowledgement

We would like to thank the following people and institutions. The VIB Bioimaging Core Leuven, in particular Nicolas Peredo, Nikky Corthout, Axelle Kerstens, and Benjamin Pavie, for the training, the facilities, and the use of IMARIS. The UZ Brussels, in particular the patients who kindly donated embryos for this research, and Yves Guns, Sabrina Vitrier, and all the other Brussels IVF personnel, for facilitating our research. The KU Leuven FACS core: Vera Dermesrobian and Barbara Moraschi, for the training, the facilities, and the help, and Susan Schlenner for leadership. The KU Leuven Genomics Core for the training and the facilities. The members of the KU Leuven Stem Cell and Development Unit, Bradley Balaton, Tijs Vanhessche, Wilhelm Bouchereau, Dylan Laurens, Jonas Degelin and Ryan Allsop, members of the VUB, the Graziano Martello lab, the Nicolas Rivron lab, the Laurent David lab and Eva Moinard, the Harunobu Kagawa lab, the Peter Rugg-Gunn lab, and the Thorold Theunissen lab, for providing help, protocols, feedback, and facilitating open science. The Statistical Consulting Group, Biomedical Sciences Leuven. Jean-Christophe Marine’s laboratory for sharing the Tapestation. The UZ Leuven, more specifically Ilse Parijs (Centre of Human Genetics) and Ronny Decorte (Department of Forensic Biomedical Sciences), for karyotyping and STR testing of the cell lines that were used, respectively. And the funding agencies FWO and KU Leuven for making this research possible. M.A.S. is supported by the Gorgas Memorial Institute for Health Studies and Fundación Sus Buenos Vecinos in Panama.

## Declaration of generative AI and AI-assisted technologies in the manuscript preparation process

During the preparation of this work, the authors used Perplexity (paid web version), ChatGPT, Claude (Sonnet 4.6, 5, unpaid), and M365 Copilot (GPT-4) to improve the language and readability of parts of the manuscript and to troubleshoot data analysis. After using these tools/services, the authors reviewed and edited the content as needed and take full responsibility for the content of the published article.

---

**Supplementary Fig. 1:**
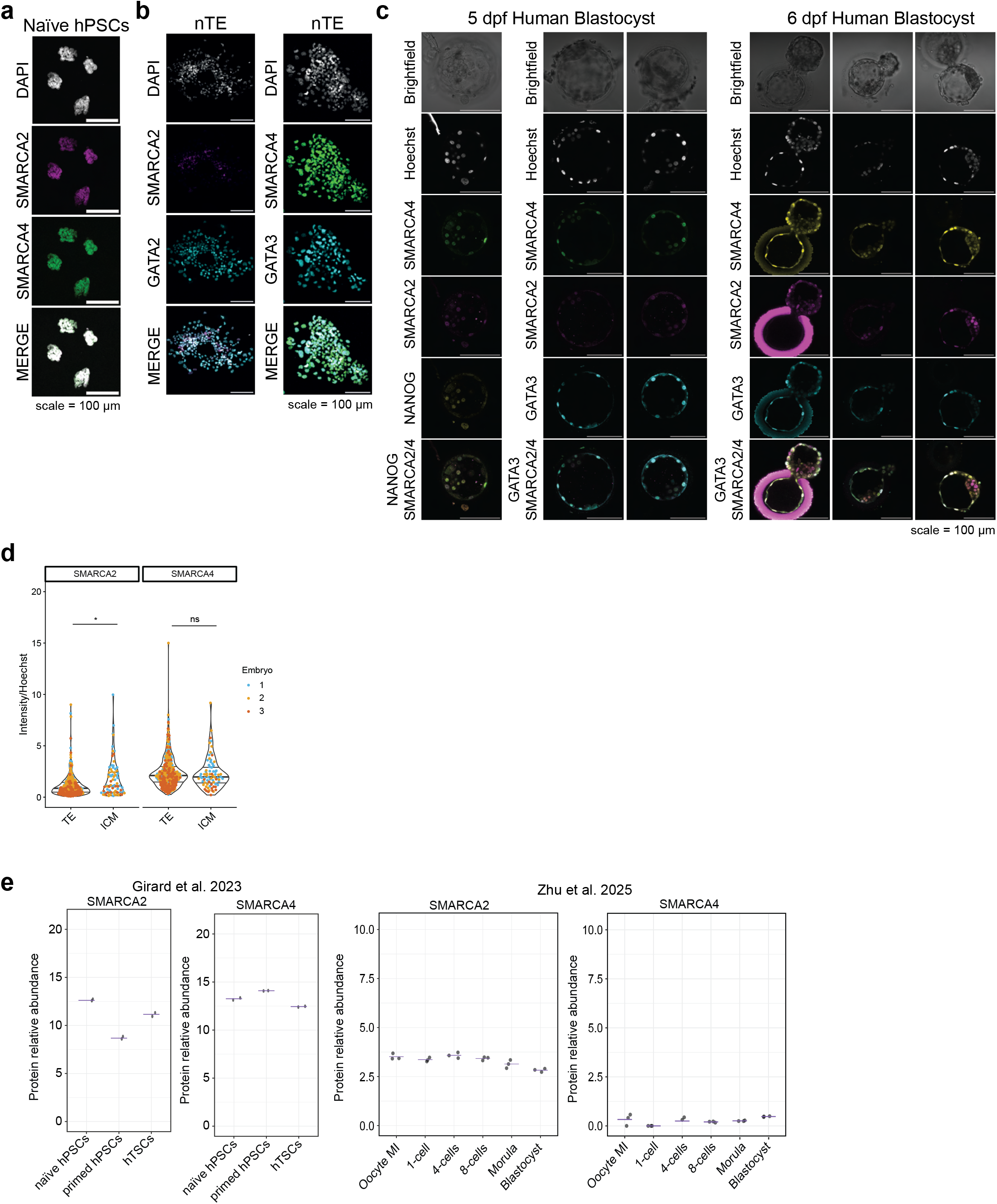
Characterisation of SWI/SNF ATPases SMARCA2 and SMARCA4. **a, b** Immunofluorescence images of naïve hPSCs **(a)** and day 3 naïve-derived trophectoderm (nTE) cells **(b)** showing SMARCA2 and SMARCA4 protein expression. Scale bar = 100 µm. **c** Immunofluorescence image of 5 days post-fertilisation (dpf) and 6 dpf human blastocysts showing SMARCA2 and SMARCA4 protein expression, together with NANOG or GATA3. Scale bar = 100 µm. N = 3 embryos. **d** Box plot showing SMARCA2 (top) and SMARCA4 (bottom) signal intensities, normalised for Hoechst intensity, of ICM and TE cells of the 6 dpf blastocysts of **Supplementary Fig. 1c**. The lines represent the 25th, 50th, and 75th percentiles. N=3 embryos. **e** Plot showing relative protein abundance of SMARCA2 and SMARCA4 in naïve hPSCs, primed hPSCs, and human TSCs (hTSCs)^52^ (left). Plots showing relative protein abundance of SMARCA2 and SMARCA4 in different stages of pre-implantation human development^53^ (right). * = p-value < 0.05; ns indicates non-significance.

**Supplementary Fig. 2:**
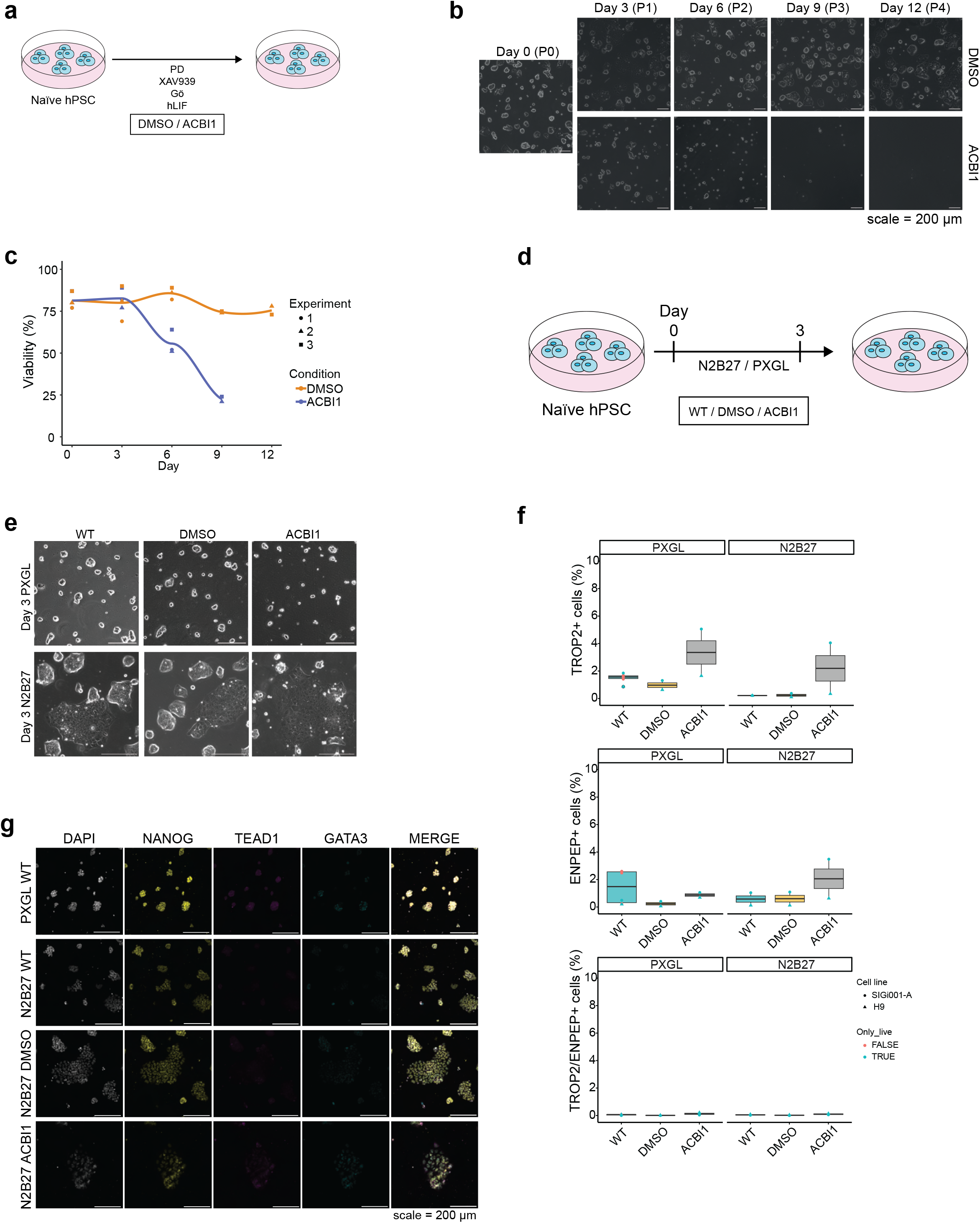
Perturbation of SWI/SNF ATPases SMARCA2 and SMARCA4. **a** Schematic overview of the experimental set-up of naïve hPSC culture in PXGL with DMSO and ACBI1. **b** Brightfield microscopic image showing H9 naïve hPSCs at day 3, 6, 9, and 12 in PXGL with DMSO or ACBI1. Scale bar = 200 µm. **c** Trypan Blue cell viability quantification at 3, 6, 9, and 12 days in PXGL with DMSO or ACBI1. Data points are shaped by experiment number. N = 3 experiments. **d** Schematic overview of the experimental set-up of naïve hPSCs cultured for 3 days in N2B27 or PXGL with no treatment (WT), DMSO, and ACBI1. **e** Brightfield microscopic image showing cells at day 3 in PXGL or N2B27 with and without DMSO and ACBI1. The scale bar represents 200 µm. **f** Box plot showing the percentage of ENPEP, TROP2, and ENPEP/TROP2 positive cells by flow cytometry. Red dots indicate flow cytometry without a cell viability dye, and blue dots indicate quantification with a cell viability dye. The shape of the dots represents the cell line used. N=2 experiments. **g** Immunofluorescence images of naïve hPSCs (HNES1) cultured in N2B27 for 3 days with and without DMSO or ACBI1 showing NANOG, GATA3, and TEAD1 protein expression. The scale bar represents 200 µm.

**Supplementary Fig. 3:**
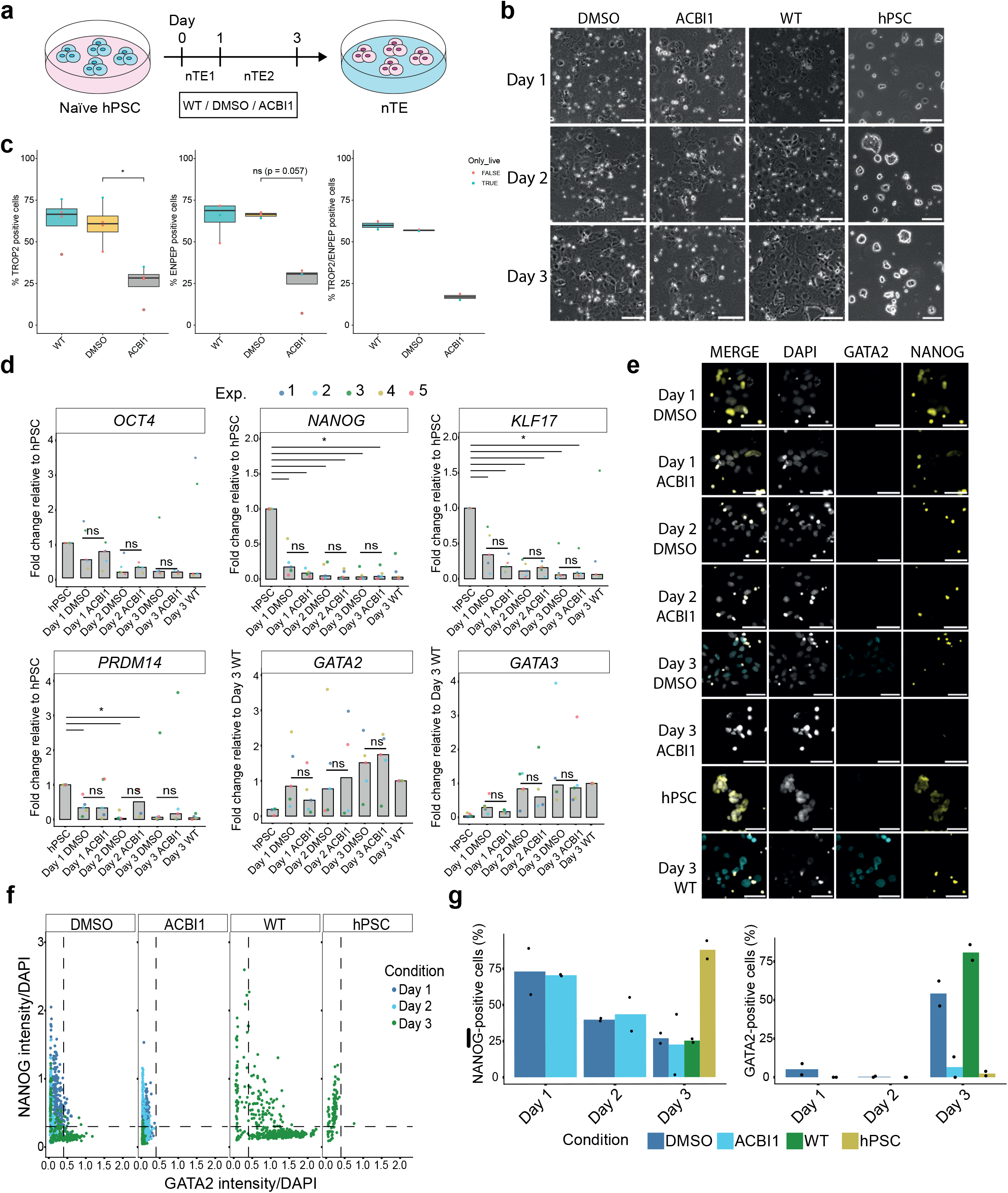
SWI/SNF perturbation during *in vitro* trophectoderm formation. **a** Schematic overview of the experimental set-up of naïve to trophectoderm conversion (nTE) with non-treated (WT), DMSO, and ACBI1. **b** Brightfield microscopic images showing cells at day 1, 2, and 3 of nTE differentiation. The scale bar represents 100 µm. **c** Box plots showing the percentage of ENPEP (N=4), TROP2 (N=4), and ENPEP/TROP2 (N=2) positive cells by flow cytometry. Red dots indicate flow cytometry without a cell viability dye, and blue dots indicate quantification with a cell viability dye. The shape of the dots represents the cell line used. **d** Bar plot showing RT-qPCR RNA expression quantification of pluripotency genes (*OCT4, NANOG, KLF17, PRDM14*) normalised to naive hPSCs, and trophectoderm genes (*GATA2 and GATA3*) normalised to day 3 nTE untreated (WT), at day 1, 2, and 3 of nTE differentiation. N = 5 experiments. **e** Immunofluorescence images of day 1, 2, and 3 of nTE differentiation showing trophectoderm marker GATA2 and epiblast marker NANOG. Scale bar = 50 µm. **f** Scatter plot showing immunofluorescence signal quantification, corrected for DAPI intensity, of trophectoderm marker GATA2 and epiblast marker NANOG at day 1, 2, and 3 of nTE differentiation. N = 2 experiments and one replicate shown. Colour indicates the time point. Dotted line indicates the cut-off for ‘positive cells’ **(Supplementary Fig. 3g)**. **g** Bar plot showing quantification of NANOG-positive and GATA2-positive cells at day 1, 2, and 3 of nTE differentiation. N = 2. Positive cells are determined based on cells above the dotted line **(Supplementary Fig. 3f)**. * = p-value < 0.05, and ns indicates non-significance.

**Supplementary Fig. 4:**
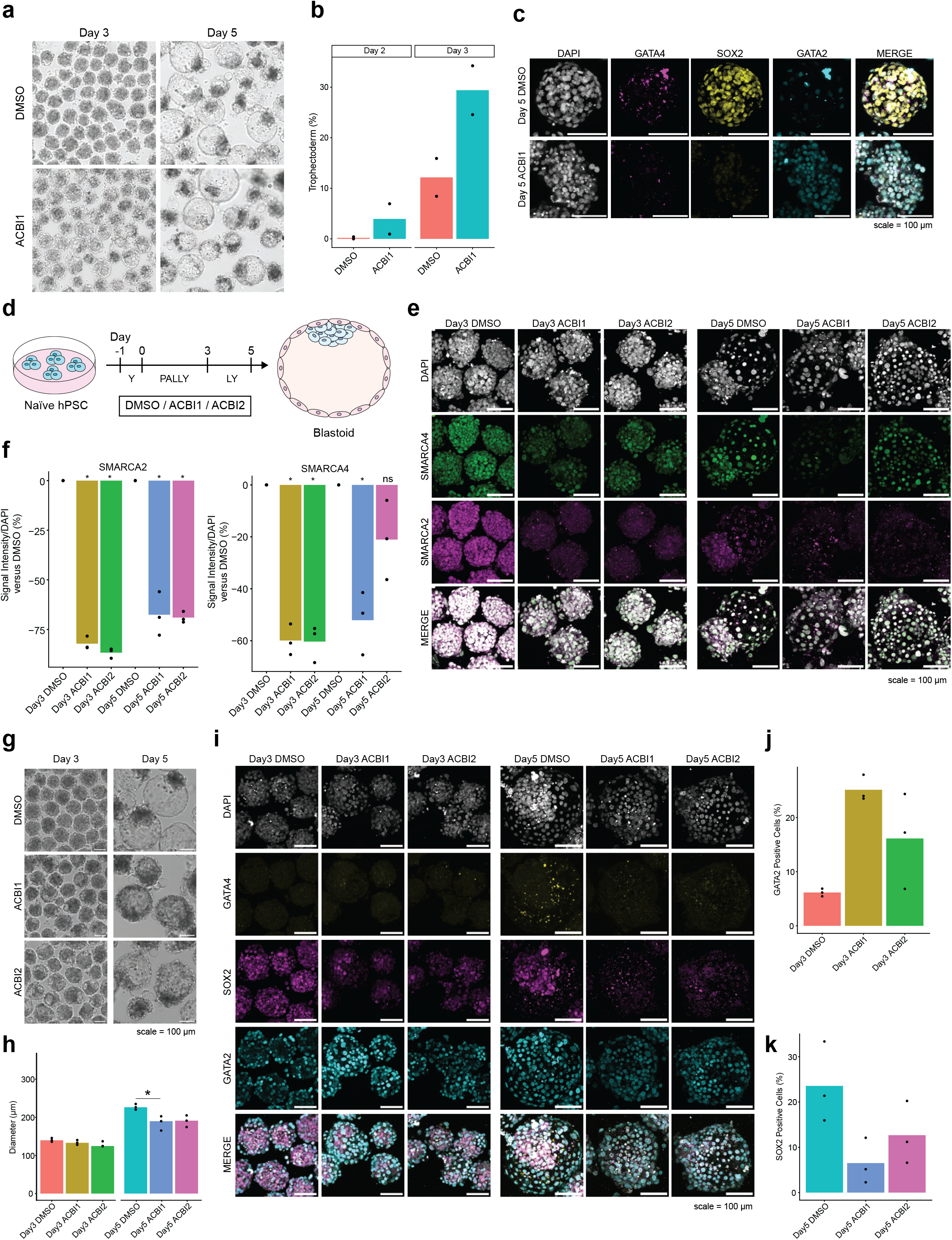
SWI/SNF perturbation during blastoid formation. **a** Brightfield microscopic image showing blastoids at day 3 and day 5 of blastoid formation. **b** Bar plot showing the quantification of trophectoderm cells in day 2 and day 3 blastoids upon ACBI1 treatment. N = 2. **c** Immunofluorescence images showing the presence of hypoblast marker GATA4, epiblast marker SOX2, and trophectoderm marker GATA2 in day 5 non-blastoid aggregates. Scale bar = 100 µm. **d** Schematic overview of the experimental setup for 50 nM ACBI1 and 5 nM ACBI2 treatment during blastoid formation. **e** Immunofluorescence images showing SMARCA2 and SMARCA4 degradation in day 3 and day 5 human blastoids upon ACBI1 and ACBI2 treatment. Scale bar = 100 µm. **f** Box plot showing the change in SMARCA2 and SMARCA4 signal intensity of ACBI1 compared to DMSO, normalised by DAPI intensity, in day 3 and day 5 blastoids. N=3 experiments. **g** Brightfield microscopic image showing blastoids at day 3 and day 5 of blastoid formation upon ACBI1 and ACBI2 treatment. Scale bar = 100 µm. **h** Bar plot showing the quantification of blastoid size over time upon ACBI1 and ACBI2 treatment on day 3 and day 5. N = 3 experiments. **i** Immunofluorescence images of day 3 and day 5 human blastoids showing trophectoderm marker GATA2, hypoblast marker GATA4, and epiblast marker SOX2. Scale bar = 100 µm. **j** Bar plot showing the quantification of GATA2-positive cells in day 3 blastoids upon ACBI1 and ACBI2 treatment. N = 3. **k** Bar plot showing the quantification of SOX2-positive cells in day 5 blastoids upon ACBI1 and ACBI2 treatment. N = 3. * = p-value < 0.05 and ns indicates non-significance.

**Supplementary Fig. 5:**
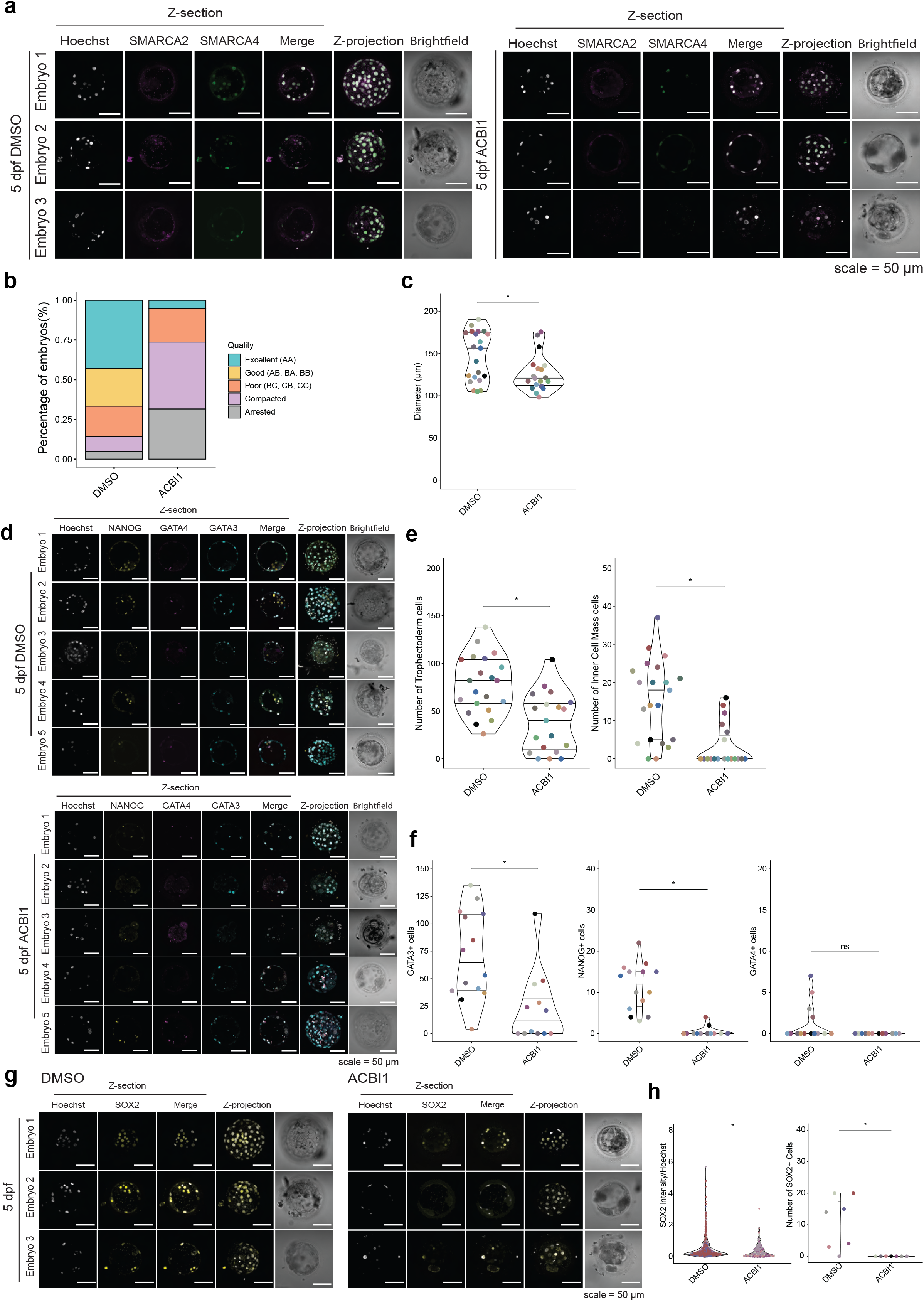
SWI/SNF ATPases preserve normal embryo development and quality. **a** Immunofluorescence images of 5 dpf blastocysts stained for Hoechst (nuclear staining), SMARCA2, and SMARCA4. Scale bar = 50 μm. **b** Stacked bar plot showing embryo quality scores. Embryos were scored based on morphological criteria^57^. Letters A-C refer to the quality of the embryo, where A is excellent, and C is poor. The first letter describes the ICM, and the second letter describes the TE. **c** Diameter without zona pellucida (ZP) of control and ACBI1-treated embryos at 5 dpf. The line represents the mean. The lines represent the 25th, 50th, and 75th percentiles. N = 21 (DMSO) and 19 (ACBI1) embryos. **d** Immunofluorescence-labelled embryos at 5 dpf for Hoechst (nuclear staining), NANOG, GATA4, and GATA3. Scale bar = 50 μm. **e** Violin plots showing the quantification of trophectoderm (TE, left) and inner cell mass (ICM, right) cell numbers. The lines represent the 25th, 50th, and 75th percentiles. N = 21 (DMSO) and 19 (ACBI1) embryos. Quantification includes embryos not stained for lineage markers. **f** Violin plot showing quantification of GATA3 (left), NANOG (middle), and GATA4 (right) positive cell numbers. The lines represent the 25th, 50th, and 75th percentiles. N = 14 (DMSO) and 12 (ACBI1) embryos. Quantification only includes embryos stained for lineage markers. **g** Representative sections and z-projections of immunofluorescent-labelled embryos at 5 dpf for Hoechst (nuclear staining) and SOX2. Scale bar = 50 μm. **h** Quantification of normalised SOX2 fluorescence intensity in embryonic cells in control and ACBI1-treated embryos. The lines represent the 25th, 50th, and 75th percentiles (left). Violin plot showing the quantification of embryo cell number for SOX2 in embryonic cells in control and ACBI1-treated embryos. The lines represent the 25th, 50th, and 75th percentiles (right). N = 7 embryos per condition. * = p-value < 0.05 and ns indicates non-significance.

**Supplementary Fig. 6:**
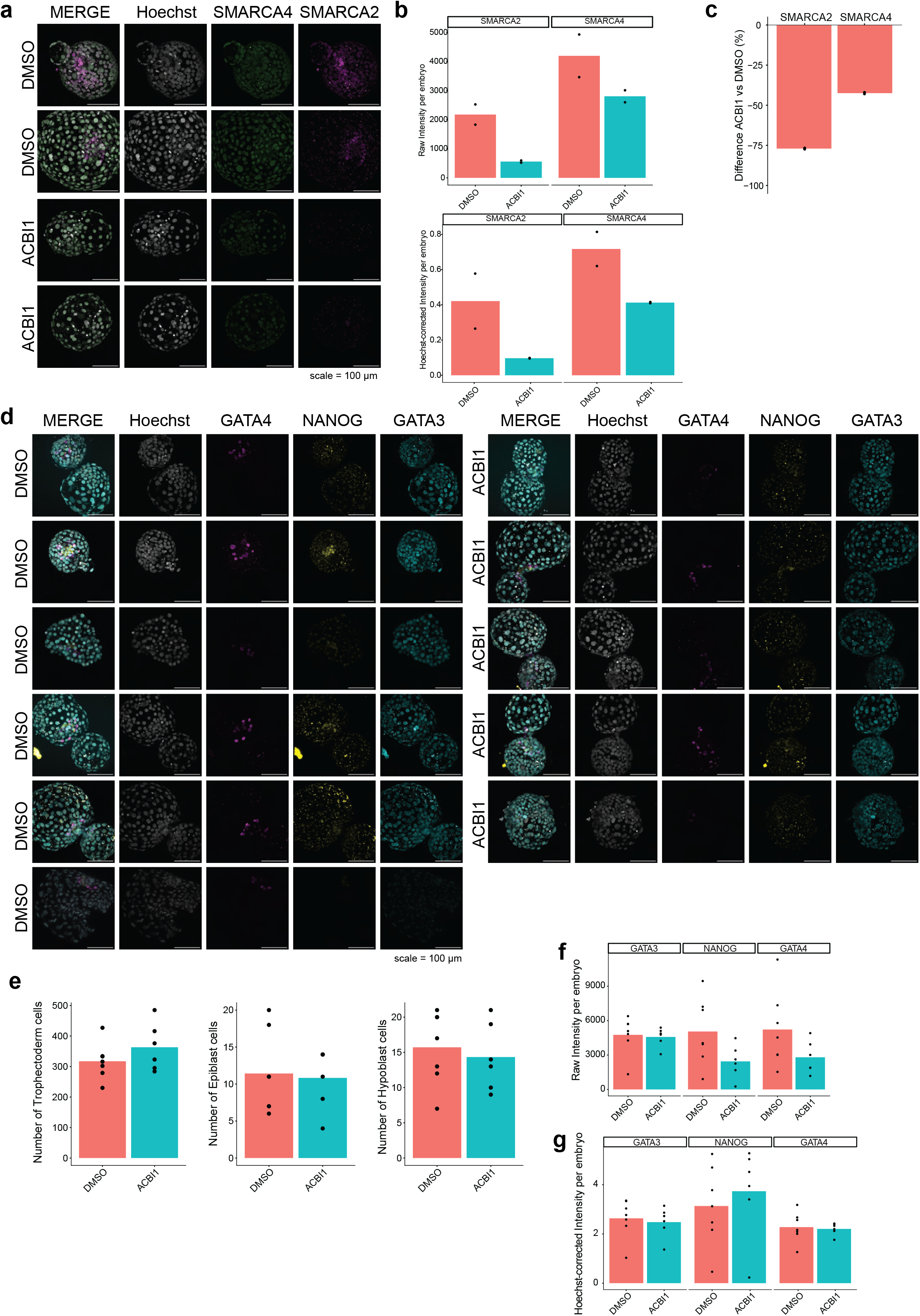
SWI/SNF ATPases perturbation during blastocyst development. **a** Immunofluorescence-labelled blastocysts at 5 dpf for Hoechst (nuclear staining), SMARCA2 and SMARCA4. Scale bar = 100 μm. **b** Bar plot showing raw (top) and Hoechst-normalised (bottom) SMARCA2 and SMARCA4 intensities. N = 2 embryos. **c** Bar plot showing relative decrease in Hoechst-normalised SMARCA2 and SMARCA4 intensities. N = 2 embryos. **d** Immunofluorescence-labelled embryos at 6 dpf for Hoechst (nuclear staining), NANOG, GATA4, and GATA3. Scale bar = 100 μm. **e** Bar plot showing quantification of trophectoderm (left), epiblast (middle), and hypoblast (right) cell numbers. N = 7 (DMSO) and 6 (ACBI1) embryos. **F** Bar plot showing Hoechst-normalised GATA3 (in TE cells), NANOG (in EPI cells), and GATA4 (in HYPO cells) intensities. N = 7 (DMSO) and 6 (ACBI1) embryos. **G** Bar plot showing raw GATA3 (in TE cells), NANOG (in EPI cells), and GATA4 (in HYPO cells) intensities. N = 7 (DMSO) and 6 (ACBI1) embryos.

**Supplementary Fig. 7:**
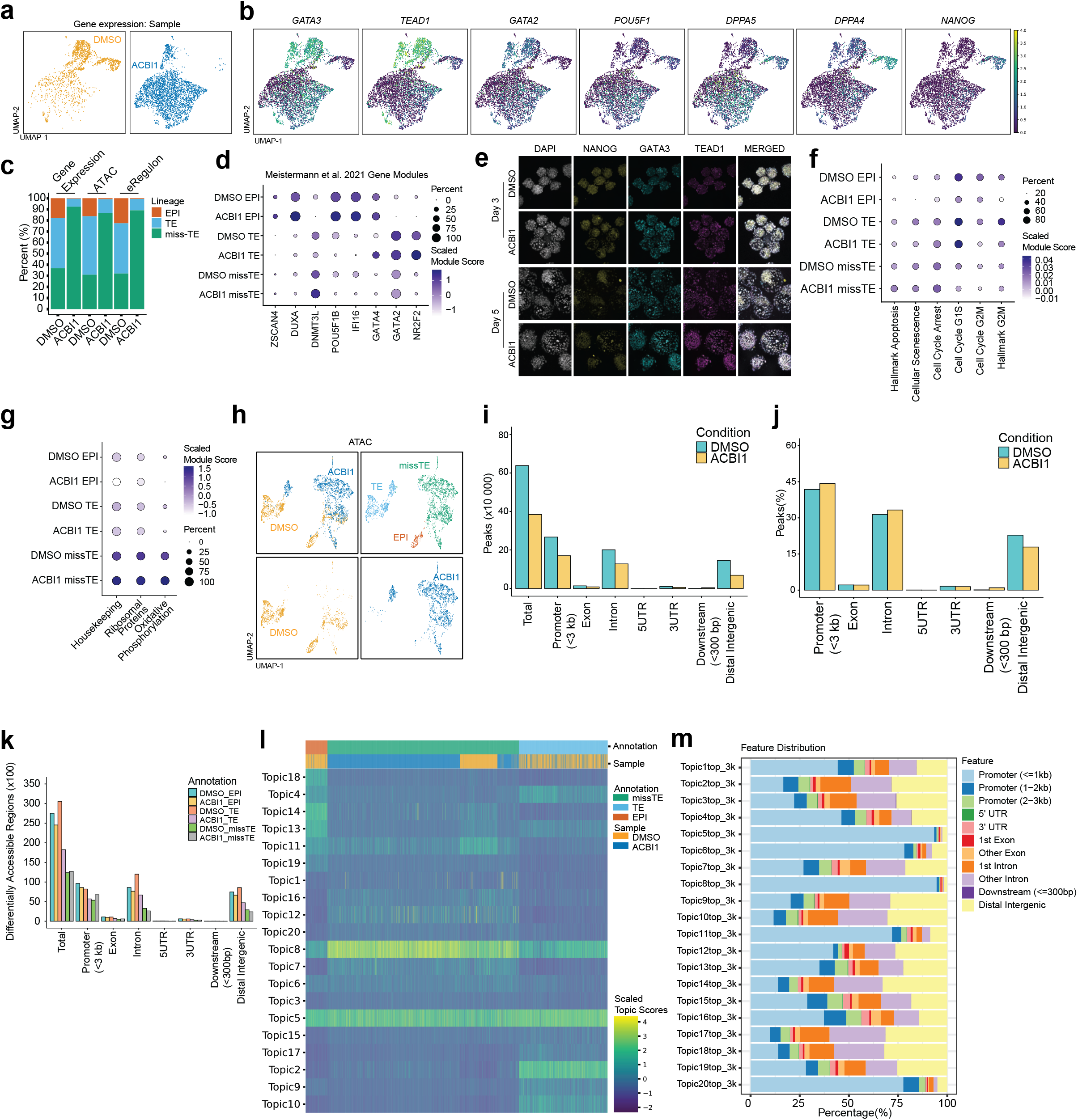
Multiomic profiling indicates that SWI/SNF ATPases facilitate the establishment of blastoid lineages through enhancer accessibility. **a** UMAP projections of the gene expression dataset, coloured and split by sample. **b** UMAP projections coloured by expression of lineage marker genes. **c** Stacked bar plot showing the percentage of cells belonging to each cluster in the gene expression, ATAC, and eRegulon datasets. **d** Dot plot showing the expression of genes in gene modules identified by Meistermann et al.^49^. Size represents the percentage of the group, and colour represents the scaled module score. **e** Immunofluorescence images showing GATA3, TEAD1, and NANOG protein expression in day 3 and day 5 human blastoids upon 50 nM ACBI1 treatment (ACBI1). **f** Dot plot showing the expression of MSigDB gene sets (M5902, M5901, M11558, M14297, M13413, M14052). Size represents the percentage of the group, and colour represents the scaled module score. **g** Dot plot showing the expression of housekeeping genes ^58^, ribosomal protein genes, and oxidative phosphorylation genes. Size represents the percentage of the group, and colour represents the scaled module score. **h** UMAP projections of the ATAC dataset coloured by sample (top left and bottom) and cell annotation (top right). **i** Bar plot showing the annotated genomic features of the peaks accumulated per sample. **j** Bar plot showing the percentage of annotated genomic features of the peaks accumulated per sample. **k** Bar plot showing the annotated genomic features of differentially accessible regions per cell type and sample. **l** Heatmap showing the scaled topic scores per annotation and sample. **m** Stacked bar plot showing the percentage of genomic features of the called topics per cell annotation and sample.

**Supplementary Fig. 8:**
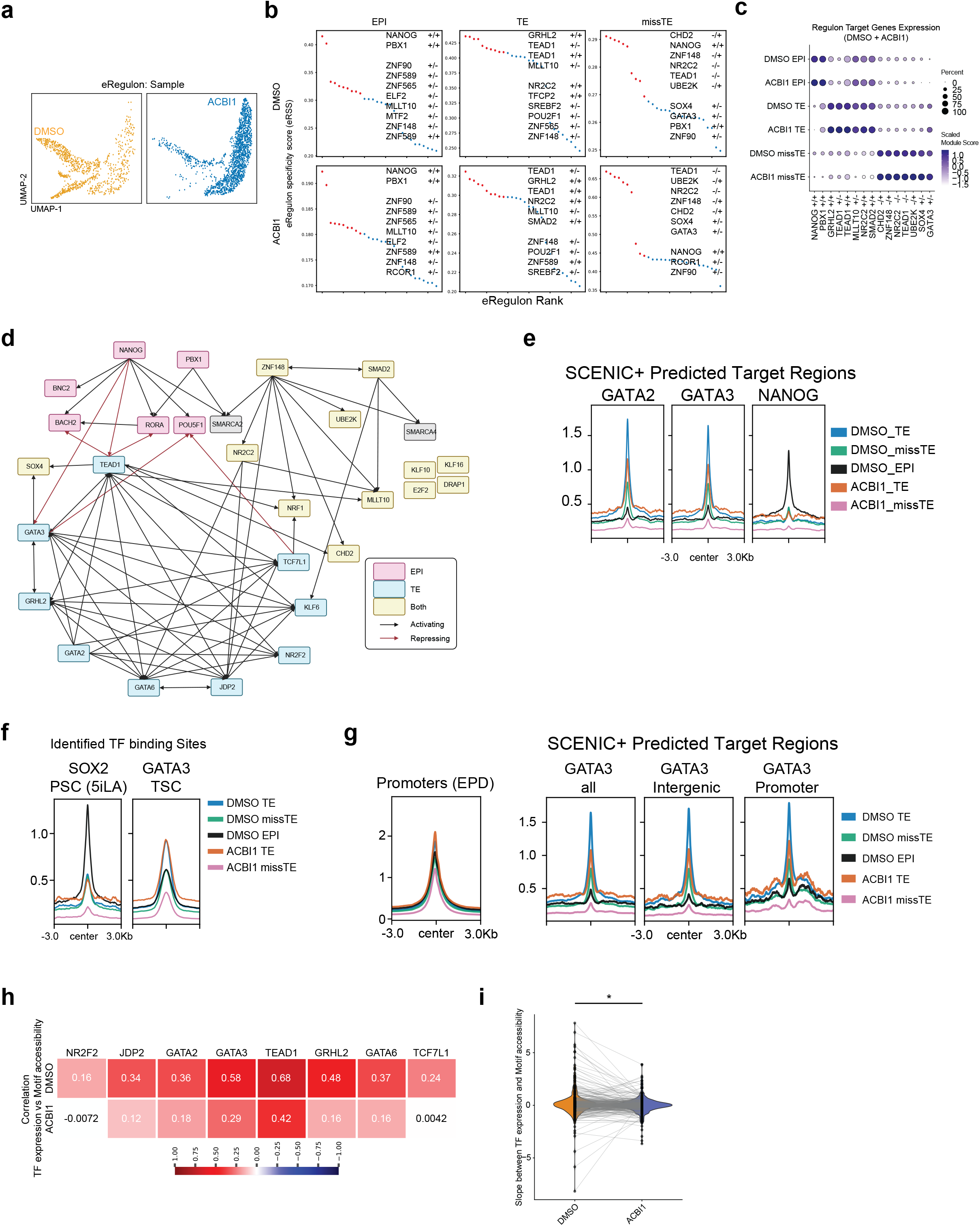
Gene regulatory network analysis in SWI/SNF-perturbed human blastoids. **a** UMAP projections of the eRegulon dataset, coloured and split by sample. **b** Plot showing eRegulon specificity score (eRSS) of eRegulons ranked per cluster. The top 10 regulons are shown in red. **c** Dot plot showing the expression of eRegulon target genes per cell type and sample. Size represents the percentage of the group, and colour represents the scaled module score. **d** Gene regulatory network showing the connection between eRegulons. Black arrow: activating. Red arrow: inhibiting. Transcription factors coloured by expression per cell type. Panel is created in BioRender (https://BioRender.com/q7xul8a) and is licensed under CC BY 4.0.. **e** Profile plot showing chromatin accessibility over SCENIC+ predicted enhancer regions targeted by GATA2, GATA3 and NANOG. **f** Profile plot showing chromatin accessibility over SOX2 in naïve hPSCs^42^ and GATA3 in TSCs^64^. **g** Profile plots showing accessibility over all promoters in the Eukaryotic Promoter Database (EPD)^88^ (left) and SCENIC+-predicted GATA3 target sites (right). **h** Heatmap showing correlation between the expression of eRegulon transcription factors and motif accessibility score by sample. **i** Violin plot showing the slope between expression and motif accessibility per transcription factor. N = 300 genes. * = p-value < 0.05.

**Supplementary Fig. 9:**
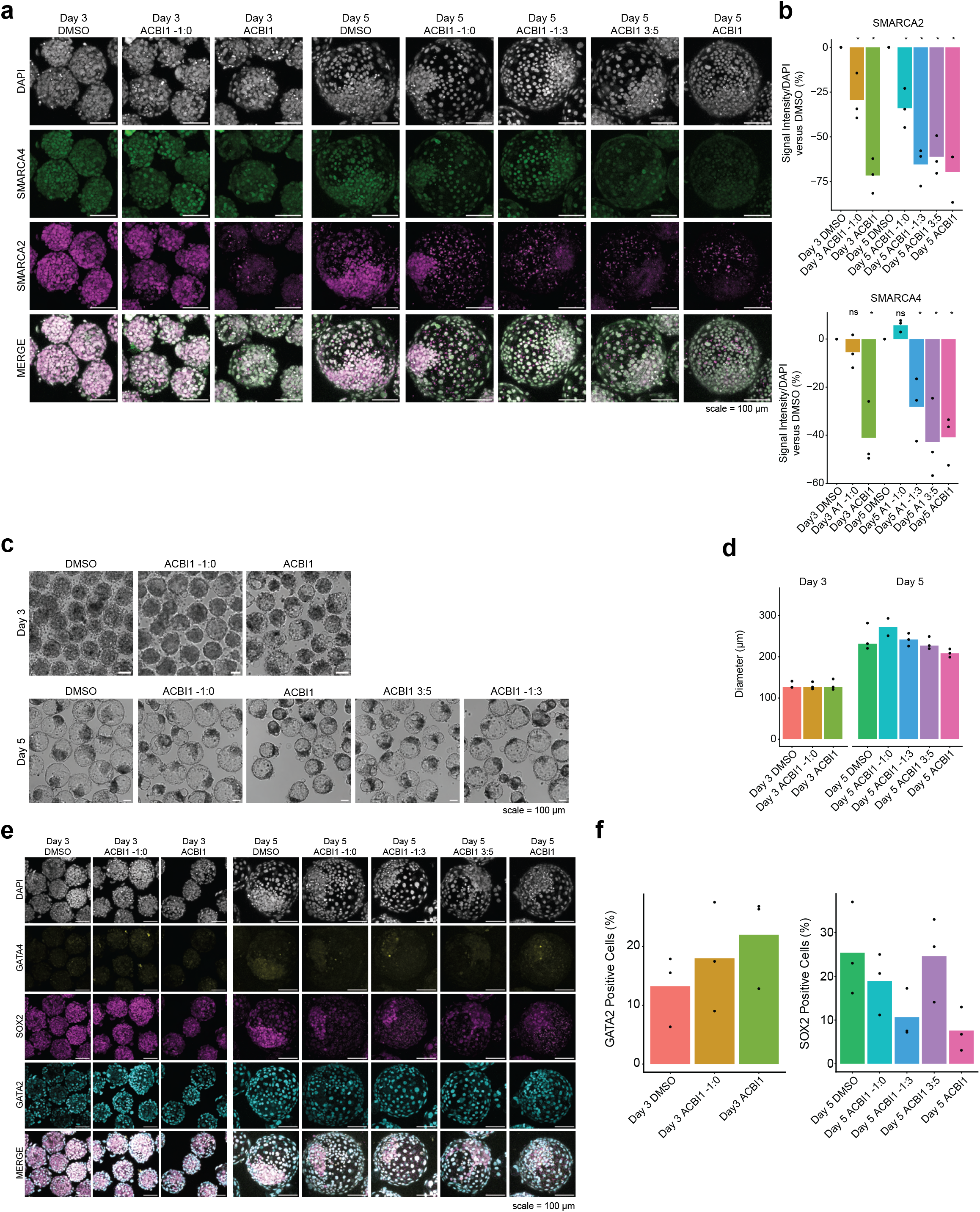
SWI/SNF ATPases regulate blastoid lineage establishment. **a** Immunofluorescence images showing SMARCA2 and SMARCA4 degradation in day 3 and day 5 human blastoids. Scale bar = 100 µm. **b** Box plot showing the change in SMARCA2 and SMARCA4 signal intensity of ACBI1 compared to DMSO, normalised by DAPI intensity, in day 3 and day 5 blastoids. N=3 experiments. **c** Brightfield microscopic image showing blastoids at day 3 and day 5 of blastoid formation. **d** Bar plot showing the quantification of blastoid size over time on day 3 and day 5. N = 3 experiments. **e** Immunofluorescence images of day 3 and day 5 human blastoids showing trophectoderm marker GATA2, hypoblast marker GATA4, and epiblast marker SOX2. Scale bar = 100 µm. **f** Bar plot showing the quantification of GATA2-positive cells in day 3 blastoids (left) and SOX2-positive cells in day 5 blastoids (right). N = 3 experiments. * = p-value < 0.05 and ns indicates non-significance.

**Supplementary Fig. 10:**
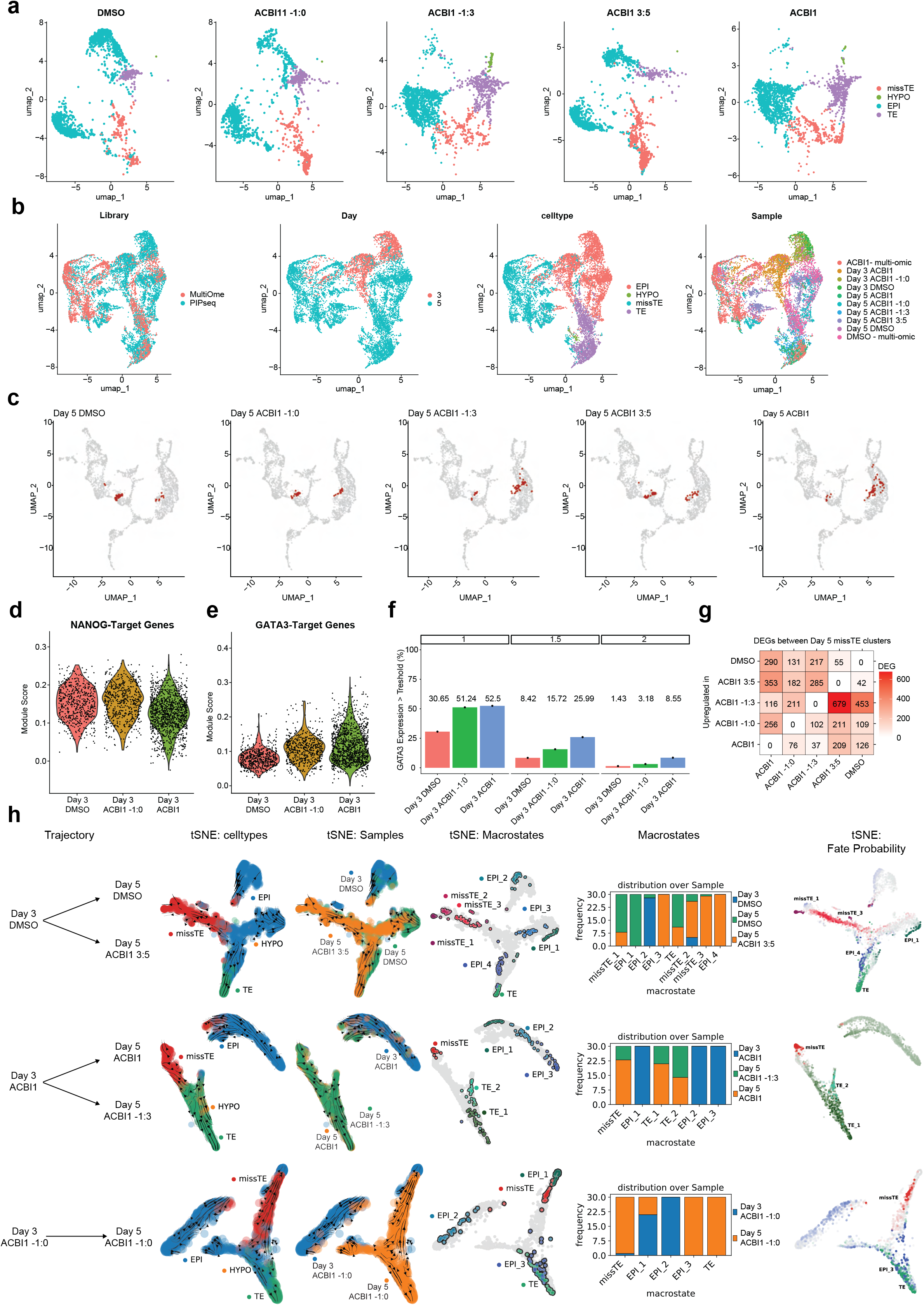
SWI/SNF ATPases affect cell lineage establishment and trajectories in blastoids. **a** UMAP projections coloured by cell type split per day 3 to day 5 trajectories. **b** UMAP projections of multiome and transcriptome integration coloured by library, day, cell type, and sample. **c** UMAP projections of gene counts of day 5 blastoids integrated with an embryo reference dataset^17^. Cells of this study are coloured in red. **d** Violin plot showing the module score of SCENIC+-identified NANOG target genes in day 3 samples. **e** Violin plot showing the module score of SCENIC+-identified GATA3 target genes in day 3 samples. **f** Bar plot showing the percentage of GATA3-positive cells in day 3 samples with expression greater than 1 (left), 1.5 (middle), and 2 (right). **g** Heatmap showing the number of upregulated genes between day 5 missTE clusters. The number shows the number of upregulated genes in the row versus the column. **h** CellRank2 pseudotime analysis showing cell trajectories between day 3 and day 5, macrostates, macrostate composition, and fate probability.

## Supplementary Information

**Supplementary Table 1:** Differential Gene expression analysis of multiomic dataset. DT: DMSO-TE. DM: DMSO-missTE. DE: DMSO-EPI. AT: ACBI1-TE. AM: ACBI1-missTE.

**Supplementary Table 2:** Blastoid Topics in BED format, top 3000 regions per topic.

**Supplementary Table 3**: All eRegulons identified by SCENIC+, their target genes, and the weight (identified in that number of iterations), related to Fig. 4 and S4

**Supplementary Table 4:** The consensus regions of the enhancers identified by SCENIC+, the eRegulon TF, their predicted target genes, enhancer metrics, and Chipseeker annotations, related to Fig. 4 and S4.

**Supplementary Table 5:** Differential Gene expression analysis of the PIPseq dataset. D3D: Day 3 DMSO. D3A0: Day 3 ACBI1 -1:0. D3A: Day 3 ACBI1. D5D: Day 5 DMSO. D5A0: Day 5 ACBI1 -1:0. D5A3: Day 5 ACBI1 -1:3. D5A5: Day 5 ACBI1 3:5. D5A: Day 5 ACBI1.

**Supplementary Table 6:** Antibodies and Primers used in this study

**Supplementary Table 7**: PIPseeker Analysis Report Metrics: Sensitivity 5

**Source Data:** Statistical Tests per figure and panel.

